# Stable working memory and perceptual representations in macaque lateral prefrontal cortex during naturalistic vision

**DOI:** 10.1101/2022.03.24.485710

**Authors:** Megan Roussy, Benjamin Corrigan, Rogelio Luna, Roberto A. Gulli, Adam J. Sachs, Lena Palaniyappan, Julio C. Martinez-Trujillo

**Author notes:** M.R, J.M.T. L.P, R.A.G, B.C, and R.L designed the research. M.R, R.L, and A.J.S performed the research. M.R, B.C, and R.A.G contributed unpublished analytic tools. M.R analyzed data. M.R created figures. M.R and J.M.T wrote the paper.

## Abstract

Primates use perceptual and mnemonic visuospatial representations to perform everyday functions. Neurons in the lateral prefrontal cortex (LPFC) have been shown to encode both of these representations during tasks where eye movements are strictly controlled and visual stimuli are reduced in complexity. This raises the question of whether perceptual and mnemonic representations encoded by LPFC neurons remain robust during naturalistic vision — in the presence of a rich visual scenery and during eye movements. Here we investigate this issue by training macaque monkeys to perform working memory and perception tasks in a visually complex virtual environment that requires navigation using a joystick and allows for free visual exploration of the scene. We recorded the activity of 3950 neurons in the LPFC (areas 8a and 9/46) of two rhesus macaques using multi-electrode arrays, and measured eye movements using video tracking. We found that navigation trajectories to target locations and eye movement behavior differed between the perception and working memory tasks suggesting that animals employed different behavioral strategies. Single neurons were tuned to target location during cue encoding and working memory delay and neural ensemble activity was predictive of the animals’ behavior. Neural decoding of target location was stable throughout the working memory delay epoch. However, neural representations of similar target locations differed between the working memory and perception tasks. These findings indicate that during naturalistic vision, LPFC neurons maintain robust and distinct neural codes for mnemonic and perceptual visuospatial representations.

**Significance Statement:** We show that LPFC neurons encode working memory and perceptual representations during a naturalistic task set in a virtual environment. We show that despite eye movement and complex visual input, neurons maintain robust working memory representations of space which are distinct from neuronal representations for perception. We further provide novel insight on the use of virtual environments to construct behavioral tasks for electrophysiological experiments.

## Introduction

Seminal lesion studies in the early 20th century demonstrated that the primate lateral prefrontal cortex (LPFC) plays a pivotal role during delayed response tasks involving maintenance of information in working memory (WM) (Baddeley, 1986; see Roussy, Mendoza-Halliday, & Martinez-Trujillo, 2021a for review). Neurons in the LPFC maintain WM representations of space (Funahashi, Bruce, & Goldman-Rakic, 1989; Goldman-Rakic, 1994; Leavitt, Mendoza-Halliday, & Martinez-Trujillo, 2017a; Constantinidis et al., 2018; Suzuki & Gottlieb, 2013; Miller, Erickson, & Desimone, 1996), as well as perceptual representations (Mendoza-Halliday, & Martinez-Trujillo, 2017; Roussy et al., 2021a). However, neurons in the LPFC are also thought to encode signals related to eye position (Bullock, et al., 2017; Hasegawa, Sawaguchi, Kubota, & Fuster, 1998; Boulay, Pieper, Leavitt, Martinez-Trujillo, & Sachs, 2016). Many of the previous studies of visual WM and perception in the LPFC that sampled neuronal activity have been conducted in conditions where gaze is constrained, and stimuli are shown on a homogenous computer screen. However, during natural vision, primates sample complex information via gaze shifts and saccades in visual scenes that contain multiple items and variable layouts. It is unclear whether perceptual and WM representations in LPFC neurons remain invariable or deteriorate under these naturalistic conditions.

One of the most universally recognized spatial WM tasks is the oculomotor delayed response (ODR) task in which animals are required to saccade to a remembered cued location (Funahashi, Bruce, & Goldman-Rakic, 1989; Leavitt, Pieper, Sachs, & Martinez-Trujillo, 2018). During the cue presentation and delay epoch of the task, animals must maintain gaze on a fixation point. Breaking fixation results in an ‘error trial’ meaning that correct performance of the task is contingent on maintaining proper eye position during the delay epoch. This intentional and task pertinent eye fixation limits the possible effect of gaze, saccades, and eye position on the measured neuronal activity. However, this strict control of eye position during memory maintenance deviates from how WM is used in naturalistic conditions. In day-to-day life, we move our eyes while using WM, yet we are able to maintain robust WM representations of locations despite those changes in eye position. It is currently unclear how unrestrained eye position in a visually complex environment may affect the ability of neurons and neuronal ensembles in the LPFC to represent perceptual and mnemonic information.

Here, we measure firing rates of neurons in the LPFC of two macaques during virtual WM and perceptual tasks while allowing the animals to freely view a rich visual environment.

We recorded the activity of 3950 neurons in the LPFC (areas 8a/9/46) (Petrides, 2005) of both animals while measuring eye position. Neuronal activity was predictive of target location during WM and perception despite changes in eye position. Eye position poorly predicted target location when compared to neuronal activity. Additionally, using linear classifiers, we found that coding of remembered and perceived targets does not generalize in LPFC neuronal populations.

## Results

### Naturalistic working memory and perception tasks

We developed a naturalistic spatial WM task using a virtual reality engine (Unreal Engine 3, UDK). The task took place in a virtual arena that allowed for free navigation using a joystick. Importantly, to simulate natural behavior, animals were permitted free visual exploration (unconstrained eye movements) during the entire trial duration. On each trial, a target was presented for 3 seconds during the cue epoch at 1 of 9 locations in the virtual arena (Fig. 1a, b). In the WM task, the target then disappeared during a 2 second delay epoch. Navigation was disabled (i.e., joystick movements did not trigger any movement in the virtual arena) during the cue and delay epochs. Subsequently, navigation was enabled, and animals were required to virtually navigate to the target location within a 10 second response period to obtain a juice reward (Fig. 1c). We also developed a perceptual version of this task in which the target remains on screen for the trial duration (Fig. 1c). We trained two rhesus monkeys (Macaca mulatta) on both virtual tasks and recorded neuronal activity during task performance using two 96-channel micro-electrode arrays (Utah Arrays) in each animal. Arrays were implanted in the left LPFC (area 8a 46/9; one on each side of the principal sulcus, anterior to the arcuate sulcus) (Fig. 1d, e) (Petrides, 2005).

**Fig. 1.**
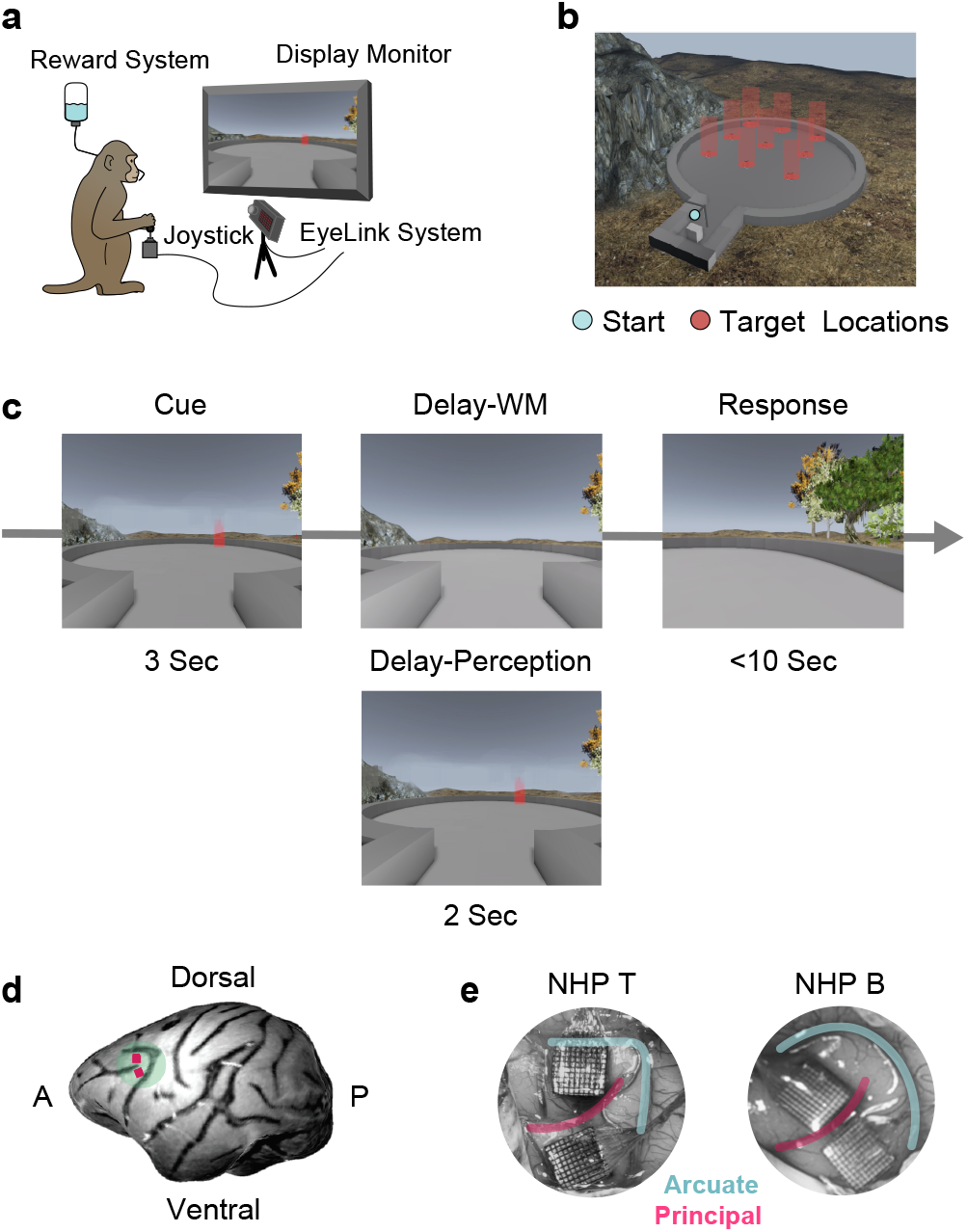
Experimental Setup. **a**. Animal in task setup with joystick, reward system, eye recording system, and monitor displayed. **b**. Overhead view of the virtual environment indicating the start location and the nine target locations. **c**. Task timeline displaying the cue, delay, and response epochs for the working memory and perception tasks. **d**. 3D modelled brain image from an MRI of NHP B with Utah array locations in the left hemisphere indicated by pink squares. **e**. Surgical images showing implanted Utah arrays in both animals.

### Task performance and animal behavior

We analyzed behavior from 20 WM sessions (12 from NHP B, 8 from NHP T) and 19 perception sessions (14 from NHP B, 5 from NHP T). Both animals performed the tasks above chance (theoretical chance 11%). Both animals performed significantly better on the perception (NHP B: Mean = 98%, NHP T: Mean = 95%) than the WM memory task (NHP B: Mean = 87%; NHP T: Mean = 57%), reflecting the increased difficulty of including a memory delay epoch (Fig. 2a). Response times for correct trials were consistent between animals and tasks (Fig. 2b).

**Fig. 2.**
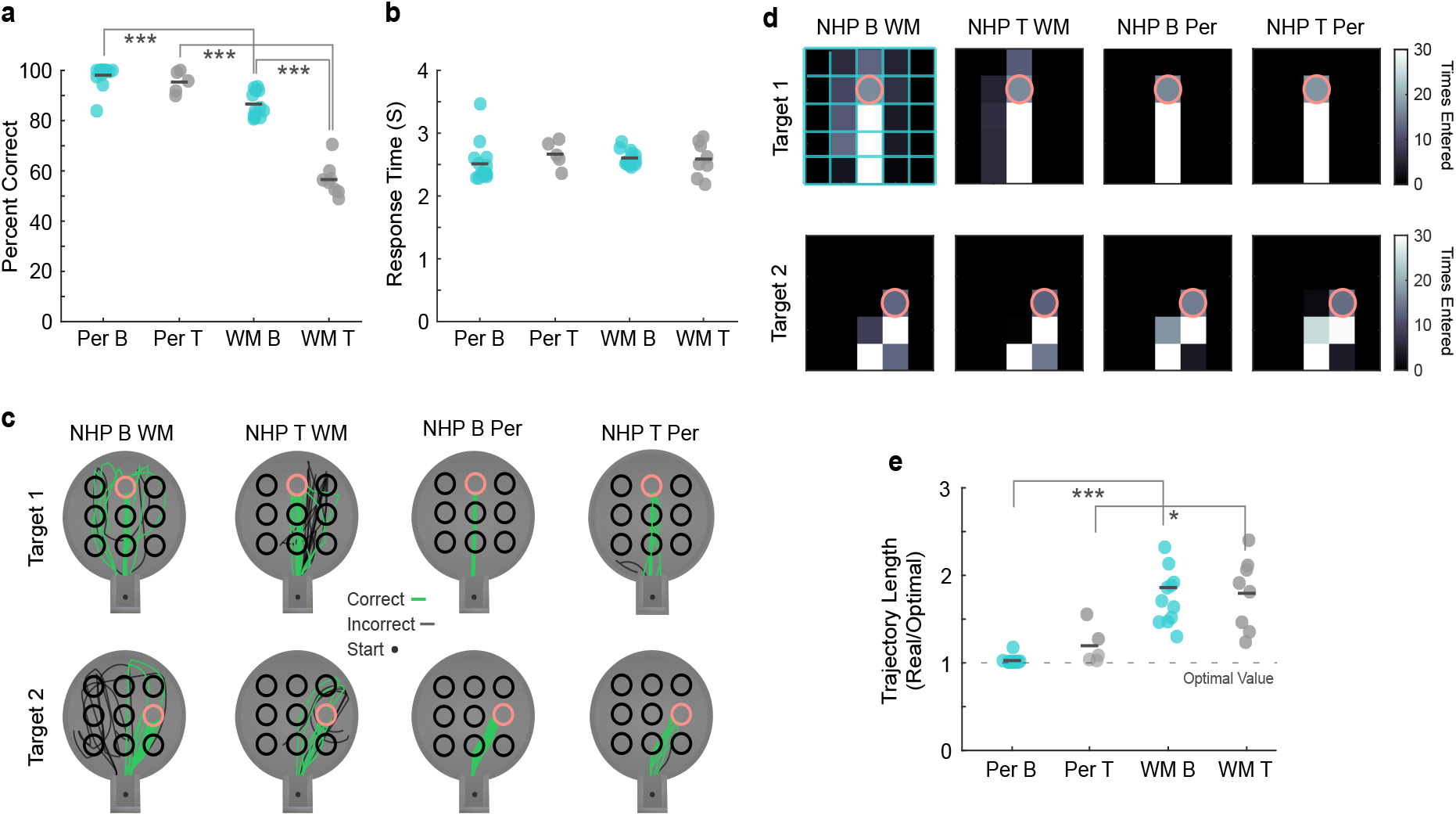
Task Behavior. **a**. Percent of correct trials for the working memory and perception tasks for each animal. Dark grey lines represent mean values and each data point represents a session. **b**. Response time for correct trials for the working memory and perception tasks for each animal. Dark grey lines represent mean values and each data point represents a session. **c**. Animal trajectories plotted for an example session and two example target locations (in pink) in which green trajectories indicate correct trials and black trajectories indicate incorrect trials. Example sessions are included for working memory and perception tasks as well as both animals. **d**. Virtual arena divided into 16 regional cells. The number of times each cell is entered (i.e. the number of trajectory points within each cell) is shown averaged over sessions for two example locations (in pink). Examples are included for working memory and perception tasks as well as both animals. **e**. Optimal trajectory measures how optimal the trajectory to correct target locations is based on path length in which a value of one, marked by the grey dashed line, reflects the shortest possible path. Optimal trajectory is plotted for the working memory and perception tasks for each animal. Dark grey lines represent median values and each data point represents a session. *p* < 0.01=*, *p* < 0.001=**, *p* < 0.0001=***.

We plotted animal trajectories to two example target locations to understand how animals were navigating in the virtual space (Fig. 2c). We divided the environment into a 16-cell grid and calculated the number of times that animals entered each cell as part of their navigation trajectory. Two example target locations averaged over all sessions are shown in Fig 2d. We next calculated the trajectory of animals in the environment in each correct trial from their starting location to the location of the target to determine how precise animals navigated towards targets. This real trajectory length was divided by the optimal trajectory length (i.e., Euclidean distance from start to target location), resulting in a measure of deviation from optimal trajectory where a value of 1 indicates that animals took the shortest possible trajectory to a target. Trajectory lengths were similar between animals during perception (NHP B: Median = 1.0; NHP T: Median = 1.1) and during WM (NHP B: Median = 1.8; NHP T: Median = 1.9). However, trajectories were more optimal during the perception task than during the WM task, indicating less precise navigation to targets during WM, when the target was not visible (Fig 2e). Overall, these results indicate that both animals used similar behavioral strategies to perform the tasks based on similar response times and trajectories.

### Eye behavior during naturalistic working memory and perception

Our virtual reality setup allowed for precise tracking of eye movement and gaze position; therefore, we measured eye movement behavior during both tasks. First, we calculated the proportion of eye position data points falling within the presentation screen. ‘Eyes off screen’ occurs when the animals close their eyes or most often, when they look away from the screen. The proportion of eye data points falling within screen boundaries differed between task epochs and between the WM and perception tasks. During WM, animals maintained eye position on the screen less during the delay epoch (Mean = 86.0%) than during the cue (Mean = 92.9%) or response epochs (Mean = 95.0%). During perception, animals maintained their eyes on the screen less during the response epoch (Mean = 81.0%) than during the delay (Mean = 89.9%) or cue epochs (Mean = 92.6%). Unlike during WM, the percentage of eye position on screen during perception cue and delay epochs showed no significant difference (Fig 3a; Data for each NHP in SFig 1a-d)

**Fig. 3.**
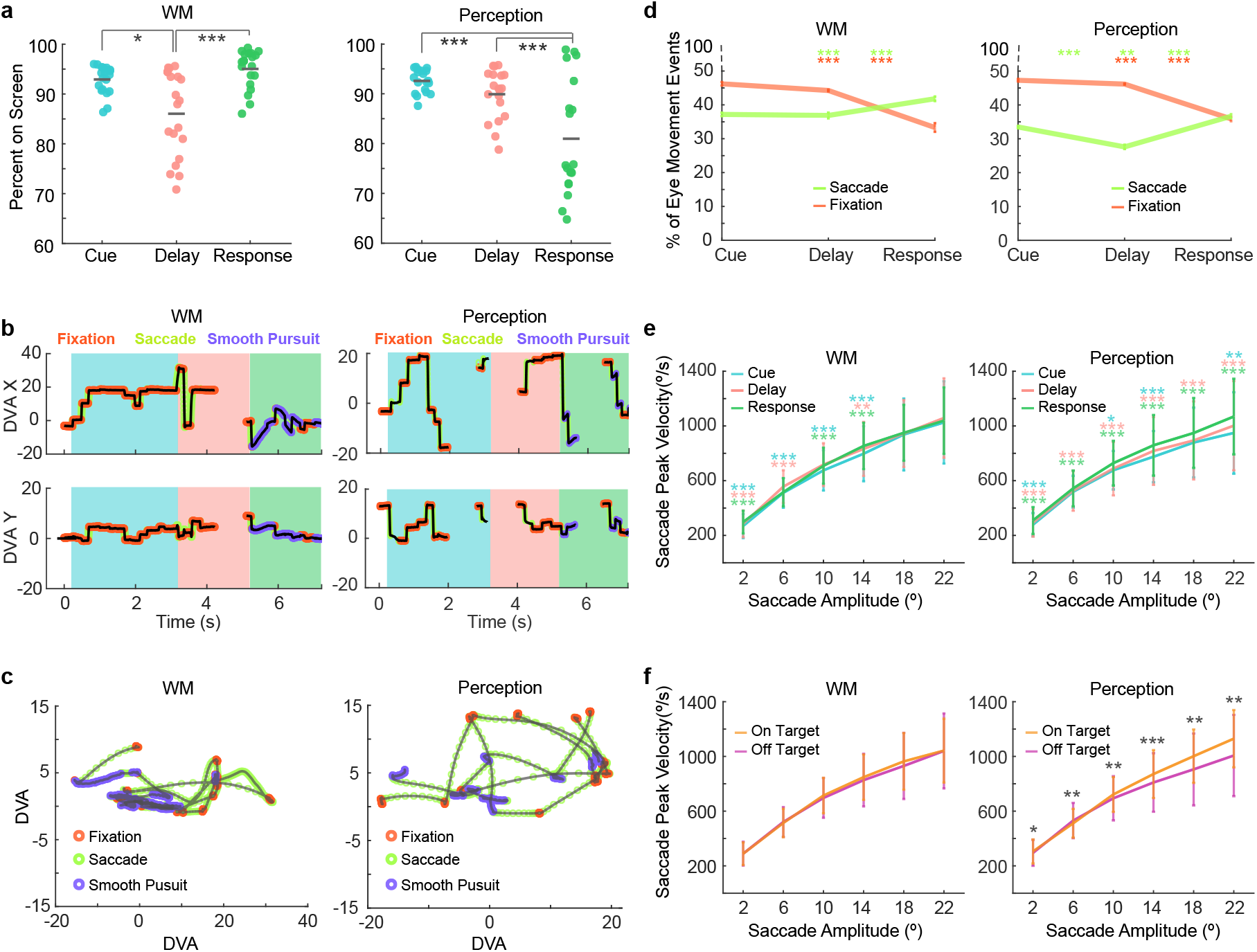
Eye Movement Behavior. **a**. Left column: the percent of eye data points that fall within the boundaries of the screen during the cue, delay, and response epochs shown for working memory sessions. Each data point represents a session. Right column: the percent of eye data points that fall within the boundaries of the screen during the cue, delay, and response epochs shown for perception sessions. Each data point represents a session. **b**. Eye traces over trial time categorized into fixations (orange), saccades (green), and smooth pursuits (purple) for an example working memory trial and an example perception trial. **c**. All eye traces categorized into fixations (orange), saccades (green), and smooth pursuits (purple) in screen coordinates for an example working memory trial and an example perception trial. **d**. Percent of eye movement events classified as fixations or saccades during the different task epochs for working memory and perception sessions. Error bars represent standard error of the mean. Asterisks on the left represent significance between the cue and delay epochs, asterisks in the middle represent significance between the cue and response epochs, and asterisks on the right represent significance between the delay and response epochs. Asterisk color corresponds to eye movement type. **e**. Main sequence for the working memory and perception task during different task epochs. Asterisks represent significance at each amplitude bin. Blue asterisks represent significance between the cue and delay epochs. Green asterisks represent significance between the cue and response epochs. Pink asterisks represent significance between the delay and response epochs. **f**. Main sequence for the working memory and perception tasks for saccades landing on and off of target location. Asterisks represent significance between on target and off target saccades at each amplitude bin. Error bars are SEM. *p* < 0.01=*, *p* < 0.001=**, *p* < 0.0001=***.

We categorized eye movement into fixations, saccades, and smooth pursuits (Corrigan, Gulli, Doucet, & Martinez-Trujillo, 2017). Example traces displaying the categorization can be found in Fig. 3b, c. We compared the proportion of eye movements that fall within each category between task epochs during perception and WM. The proportion of eye movements classified as fixations significantly differed between trial epochs and between WM and perception tasks (Fig. 3d; Data for each NHP in SFig 1e-h). During WM, animals made the most fixations during the cue epoch with fewer made during the delay and response epochs (Cue: Mean = 46.2%; Delay: Mean = 44.3%; Response: Mean = 33.3%). During perception, animals also fixated the least during the response epoch with more fixations made during the cue and delay epochs (Cue: Mean = 47.3%; Delay: Mean = 46.2%; Response: Mean = 35.9%).

The proportion of eye movements classified as saccades significantly differed between trial epochs and between WM and perception tasks. During WM, the proportion of saccades was highest in the response epoch with fewer occurring in the cue epoch and fewest during the delay epoch (Cue: Mean = 37.2%; Delay: Mean = 36.9%; Response: Mean = 41.8%) (Fig. 3d, left panel, Data for each NHP in SFig. 1e, f). During perception, animals also made the highest proportion of saccades during the response epoch (Mean = 36.7%) with fewer occurring during the cue (Mean = 33.5%) and delay epochs (Mean = 27.6%) (Fig. 3d, right panel, Data for each NHP in SFig. 1g, h). Between WM and perception response periods, there was a larger proportion of smooth pursuits during perception (Mean = 30.4%) than during WM (Mean = 28.1%). The latter may be linked to the presence of the target during perception but not during WM.

Between the WM and perception delay epochs, there was a larger proportion of eye movements classified as saccades that occur in WM (Fig. 3d; Data for each NHP in SFig. 1e-h). There were also more eye movements on screen time (Fig. 3a, b; Data for each NHP in SFig. 1a-d) and a higher proportion of saccades that occur during the WM response epoch compared to the perception response epoch (Fig. 3d; Data for each NHP in SFig. 1e-h).

To further explore saccadic activity, we calculated the main sequence, reflecting the relationship between saccade peak velocity and amplitude (see methods) (Fig. 3e; Data for each NHP in SFig. 2a-d). Saccade velocity was significantly different (higher peak velocities as a function of saccade amplitude) in the response epoch compared to the cue and delay epochs during perception for all amplitude bins (t-test, p < 0.05, effect size > 0.2). The increased velocity of saccades during perception response may reflect the use of saccades to track the target during navigation which does not occur during WM when the targets were no longer present (Fig. 3e; Data for each NHP in SFig. 2a-d). It may also signify an increase in arousal during navigation, which would be more demanding than the other task epochs. We also compared the main sequences between saccades that land on target and off target during the delay epoch (Fig. 3f; Data for each NHP in SFig. 2e-h). We found that on-target saccades resulted in larger peak velocities; however, these differences were more pronounced and were only significant during the perception task (WM: t-test, p > 0.05; Perception, t-test, 6 bins, p < 0.05). Therefore, saccades that land on target versus those that land off target show a greater difference when the target was physically present compared to when it was removed during the WM delay.

These behavioral results indicate a difference in animal behavior during different task epochs and between WM and perception. In particular, less time spent looking onscreen during the delay epoch of the WM task combined with fewer fixations, and no significant differences in saccade amplitude to targets compared to off targets suggests that animals were less focused on the target location - likely due to its removal. It is possible that animals searched for landmarks that could serve as references for the target location. Decreased fixation and increased number of saccades during the response epoch as well as an increase in saccade peak velocity may suggest a similar strategy as well as reflect the dynamic nature of the task’s response epoch in which the visual environment changes as the animal changes position in the arena.

### Spatial selectivity in single neurons

We recorded the activity of 3950 units between the dorsally (1992 units) and ventrally (1958 units) placed multi-electrode arrays. Many units in this sample displayed delay activity. Fig. 4ab shows activity patterns of two neurons that selectivity increased their activity during the delay epoch for preferred target locations. Tuning for target location was identified in the population for cue and delay epochs (Cue: Ventral: Mean = 22%, Dorsal: Mean = 16%; Delay: Ventral: Mean = 14%, Dorsal: Mean = 12%) and many neurons were tuned during both the cue and delay epochs (Ventral: Mean = 37%; Dorsal: Mean = 48%) (Fig. 4cd; Data for each NHP in SFig. 3ab).

**Fig. 4.**
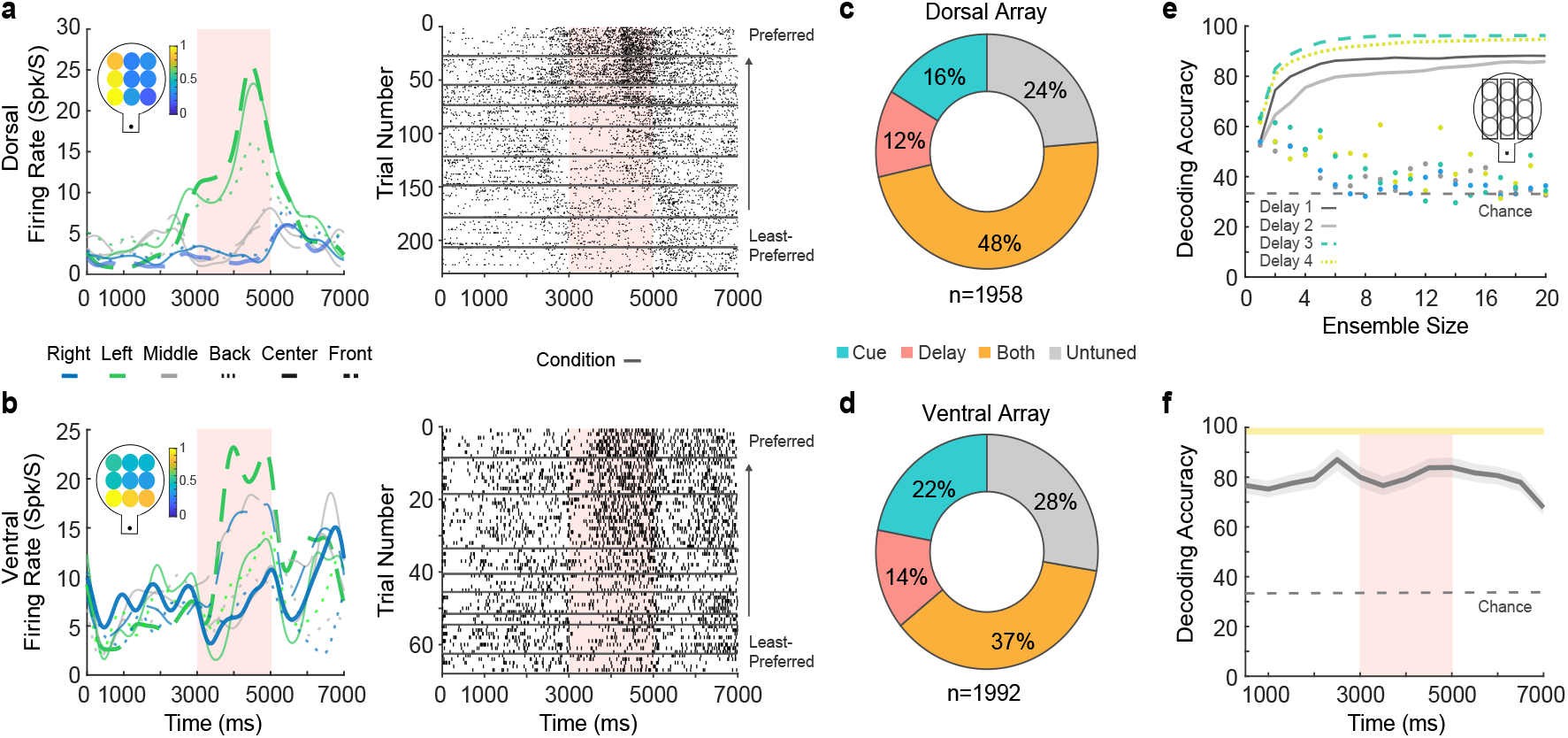
Neural Coding for Remembered Locations. **a**. Neural activity for an example neuron recorded by the dorsally located electrode array. The spike density function in the left panel displays the neuron’s activity over trial time for the nine target locations. The inlet displays normalized firing rate for all target locations. The right panel displays a raster for the same neuron in which trials are sorted by preferred to least-preferred target locations. The delay period is indicated by the salmon-colored column. **b**. Neural activity for an example neuron recorded by the ventrally located electrode array. **c**. The proportion of tuned neurons during the cue (blue), delay (pink/salmon) and during both the cue and delay epochs (orange) recorded in the dorsally located array. **d**. The proportion of tuned neurons recorded in the ventrally located array. **e**. Decoding accuracy for one example session during the delay period divided into four 500 ms temporal segments using neural ensembles of different sizes. The inlet illustrates grouping of targets into three classes. The dashed grey line represents chance decoding performance. Dots represent the decoding accuracy of individual neurons. **f**. Median decoding accuracy over trial time. The salmon-colored column represents the delay period and the grey dashed line represents chance decoding. The yellow bars on top of the figure represent significance from chance performance for each time window (t-test, *p* < 0.05). Error bars are SEM.

To determine how much information about the remembered target locations was contained in the population of neurons, we used a linear classifier (SVM - Support Vector Machine) to decode target location from neuronal firing rates within 500 ms time bins. We used a best ensemble method in which the most informative unit was found and was paired with all other neurons in the population until the best pair was found. The best pair was grouped with all neurons in the population until the best trio was found. This process was continued until the ensemble contains 20 neurons (Leavitt, Pieper, Sachs, & Martinez-Trujillo, 2017b). In order to achieve a sample size required for training and testing the classifier for all sessions, we combined trials from all targets located on the right, left, and center of the environment so decoding was performed using three classes. An example session in Fig. 4e shows decoding accuracy for different ensemble sizes during the delay epoch divided into four 500 ms time segments. Decoding accuracy over time was above chance (33.33%) for all time windows, ranging from 68% during the last 500 ms of the included response epoch to 87% towards the end of the cue epoch (Fig. 4f; Data for all sessions in SFig. 3c). The decoding accuracy was consistent over the delay epoch using both neural ensembles (Fig. 4f) and full populations of simultaneously recorded neurons (SFig. 5i), indicating robust information content for remembered locations during our naturalistic task. Decoding was also performed in 13 sessions using 9 target locations, resulting in comparable decoding accuracy (Fig 6a, b).

**Fig. 5.**
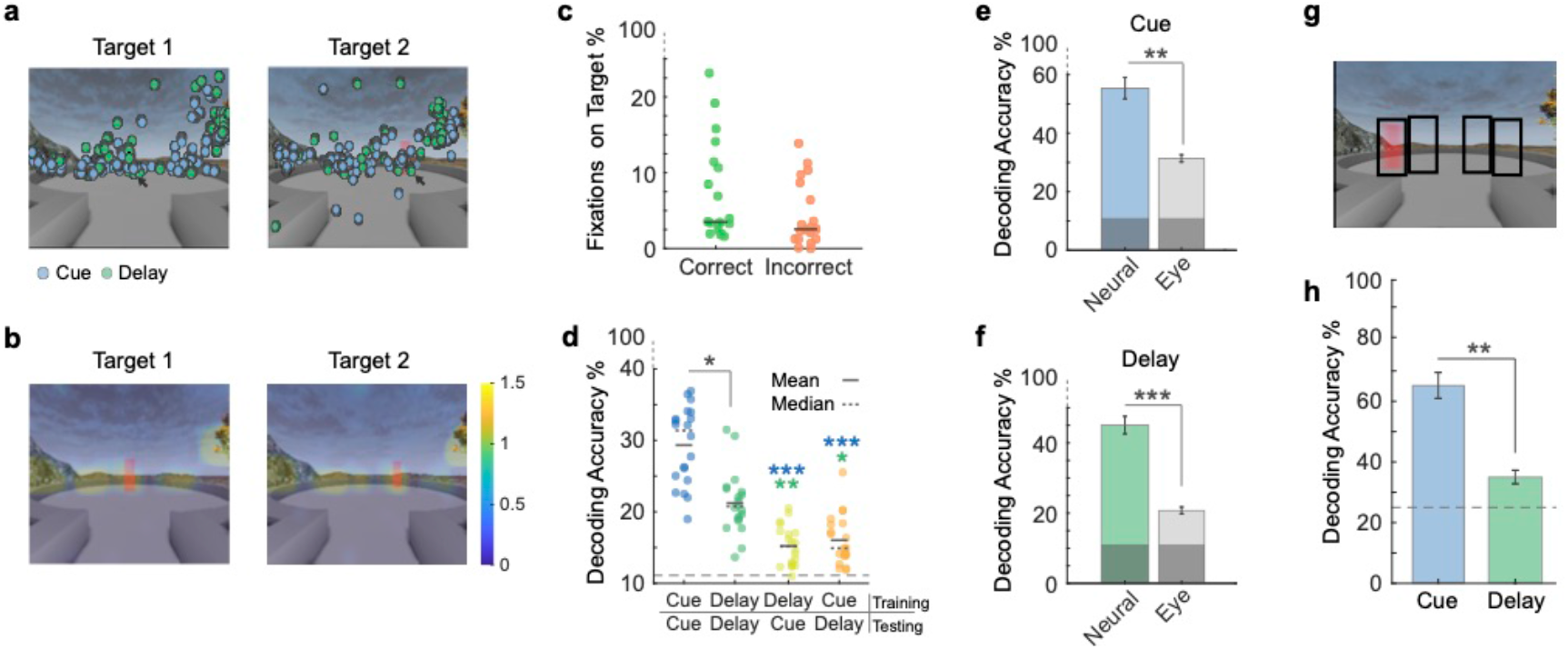
Fixations on Screen and Target Locations. **a**. All fixations for an example session plotted on screen for the cue period (blue) and delay period (green) for two example target locations. **b**. Heat maps averaged over working memory sessions showing fixation locations on screen during the delay period for two example target locations. **c**. The percentage of total fixations that fall within the target location during the delay period for correct and incorrect trials. Dark grey lines represent median values and each data point represents a session. **d**. Decoding accuracy for predicting target location from the location of eye fixations on screen during the cue and delay epochs. Classifiers are trained on the first epoch listed in the x-axis label and tested on the second epoch. The dashed line represents chance decoding accuracy. **e**. Median decoding accuracy for predicting target location based on eye position data and neural population data during the cue period. Grey regions on bars represent chance performance. **f**. Median decoding accuracy for predicting target location based on eye position data and neural population data during the delay period. Grey regions on bars represent chance performance. **g**. Outlined regions of the screen encompassing four target locations that are separable onscreen. **h**. Median decoding accuracy predicting eye position within the outlined regions shown in panel ‘g’ using neural population data during fixation periods during the cue and delay epochs. The grey dashed line represents chance decoding accuracy. Error bars are SEM. *p* < 0.01=*, *p* < 0.001=**, *p* < 0.0001=***.

**Fig. 6.**
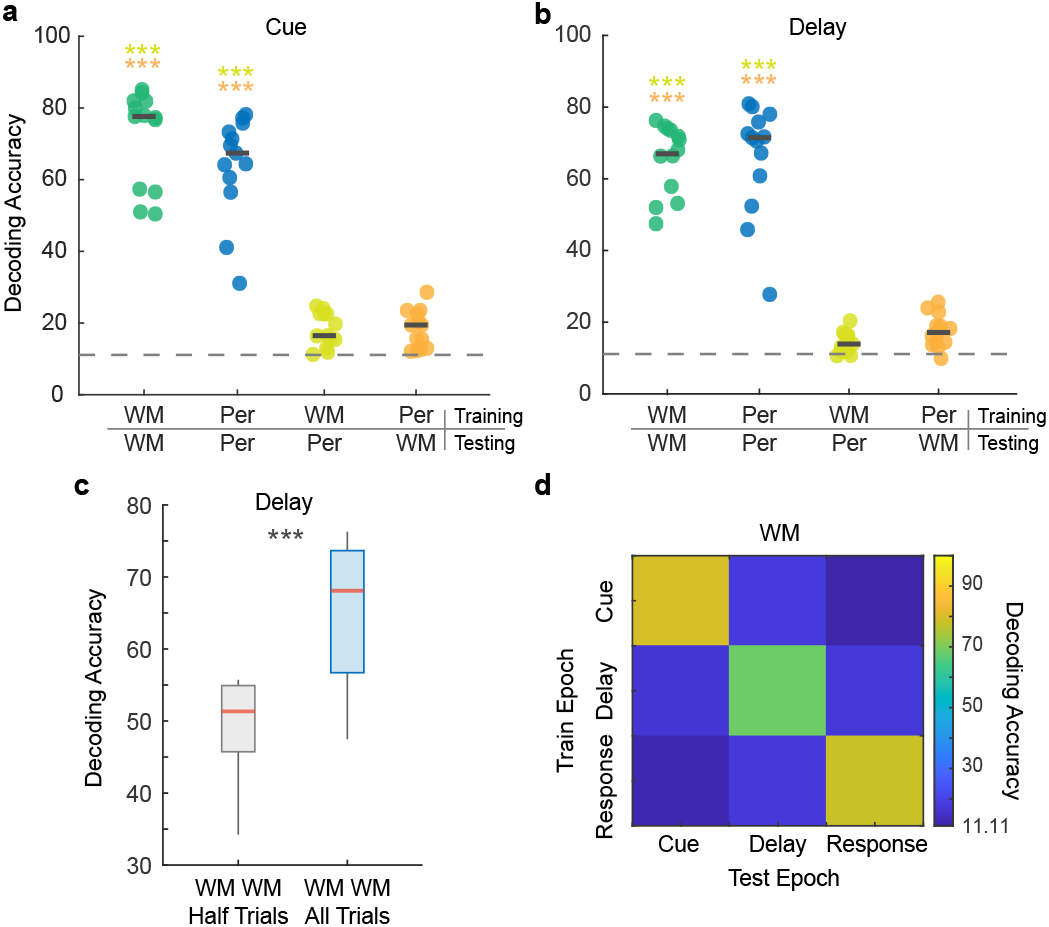
Neural Coding for Working Memory and Perception. **a**. Decoding accuracy for predicting target location for the perception and working memory tasks during the cue epoch. Classifiers are trained on the task that appears first in the x-axis label and tested on the task that appears second. Asterisk color represents significant differences with the condition of that color. Dark grey lines represent median values. The dashed grey line represents chance decoding. **b**. Decoding accuracy for predicting target location for the perception and working memory tasks during the delay epoch. **c**. Decoding accuracy for the working memory task during the delay epoch using all trials for classifier training and testing or training on half of the trials and testing on half of the trials. The red lines represent median values and the bottom and top edges of the box indicate the 25th and 75th percentiles. The whiskers extend to non-outlier data points (within 1.5 std). **d**. Cross-epoch median decoding accuracy for the working memory task. p < 0.01=*, *p* < 0.001=**, *p* < 0.0001=***.

### Fixation on target location

One potential issue in allowing for natural eye movements is that animals could maintain their gaze on the empty cued location during the delay or visually ‘rehearse’ their movement plan. We explored this possibility by analyzing gaze behavior on the targets. We plotted all fixation points on the screen for one session for two example target locations (Fig. 5a). Fixation points span the horizontal extent of the screen (constitutes the task relevant area). Fig. 5b shows heat maps of fixation locations averaged over all sessions for two example target locations during the delay epoch. Gaze was not limited to the location in which the target was presented. It was also directed to non-target stimuli in the environment such as the tree, as would occur in naturalistic contexts.

To examine if increased fixation on the cued target location was used as a behavioral strategy to improve performance, we calculated the percent of fixations falling within the bounds of the target’s location. Overall, the percentage of fixations on the target location was very low during the delay epoch (Median = 3%). There was no significant difference between correct and incorrect trials suggesting that increased fixation on cued target locations during the delay epoch may not be an effective strategy in correctly performed trials. (Correct: Median = 3.5%; Incorrect: Median = 2.6%; Wilcoxon Rank-Sum, p > 0.05) (Fig. 5c; SFig. 4a).

To determine how predictive fixation location was of target location, we divided the screen into 16 cells and calculated the number of fixation points that fell within each cell during the cue and delay epochs. We trained a SVM classifier with a linear kernel to predict which of the nine target locations was presented based on where on screen the animal was fixating. The classifier performed above chance (11.11%) during both epochs but performed significantly better during the cue epoch (Median Decoding Accuracy = 31.4%) compared to the delay epoch (Median Decoding Accuracy = 20.8%), suggesting reduced patterns of target specific fixation during the delay (Fig. 5d; SFig. 4b). To determine whether eye fixation was similar between cue and delay epochs of the WM task, we trained classifiers using eye fixation position during the cue epoch and tested the classifiers using eye fixation positions from the delay epoch. We similarly trained classifiers on delay data and tested them on cue data. Decoding accuracy was close to chance level (11.11%) when classifiers were cross-trained between epochs of the WM task and it was significantly lower than training and testing on congruent epochs (Fig. 5d; SFig. 4b). This shows that the position of fixations (i.e., gaze position) were different between the cue and delay epochs during the WM task.

We compared these results with a classifier that uses neuronal firing rates with the same number of features as the eye position classifier (16 neuron ensemble). The population of neurons contained more information about target location than gaze position in both the cue (Median Decoding Accuracy = 55.4%), and delay epochs (Median Decoding Accuracy = 45.1%) (Fig. 5e, f; SFig. 4c-f). This suggests that a population of 16 neurons encodes more information about remembered locations than the pattern of gaze positions.

### Decoding of gaze position from neural activity in LPFC neurons

Previous studies have shown that LPFC neurons encode information related to eye movements (Bullock, Pieper, Sachs, & Martinez-Trujillo, 2017). To corroborate these findings we examined whether neuronal activity in our sample of LPFC neurons contained information about the animals gaze position. We selected four targets shown in Fig. 5g that were non-overlapping on the screen and measured neuronal firing rates while animals fixated each one of the target locations. We used SVM classification and found that we could decode the gaze position from neural activity. The decoding accuracy was significantly higher during the cue epoch (Median Decoding Accuracy = 65.4%) of the WM task compared to the delay epoch (Median Decoding Accuracy = 35.0%) (Fig. 5h; SFig. 4g, h), suggesting that more information was available to the neuronal population when animals fixate on a target that was present on screen compared to when the target was no longer present. Indeed, the decoding accuracy during the delay epoch was close to chance (25%) suggesting that firing rates during fixation in the delay period carried little information about the remembered target location. One possible explanation for this finding is that decoding during the cue epoch may have been dominated by visual responses to the target. During the delay epoch, when no visual cue was present, eye position contributes poorly to decoding. These findings suggest that eye position signals do not primarily contribute to the ability of LPFC neurons to encode WM representations in complex and dynamic environments.

### Separation between coding for working memory and perception

Unlike the WM task, during the perception task, the target was accessible throughout the trial. Thus, it is possible that some neurons respond to the target only when it was present in the perception task (perceptual neurons) and some neurons are only active during the delay period of the WM task (mnemonic neurons) (Mendoza-Halliday et al., 2017; Roussy et al., 2021a). Therefore, we hypothesized that neural population activity profiles differ during the perception and WM tasks. To test this hypothesis, we collected neuronal data from 13 sessions in which animals performed both the WM and perception tasks. The same population of simultaneously active neurons were recorded during both tasks during these sessions. This allowed us to use SVM classification to cross-train neural data between WM and perception to predict the 9 target locations. We specifically tested the prediction that SVM classifiers trained in one task will not generalize the performance to the other task.

During the cue and delay epochs, decoding performance was similar between WM and perception when classifiers were trained and tested on the same task (Fig. 6a, b; SFig. 5a-d). The same population of neurons can maintain similar amounts of information about target location whether targets remain on screen (perception) or disappear (WM) (Perception: Median Decoding Accuracy = 71.5%; WM: Median Decoding Accuracy = 68.1%). Although the same neurons were recorded during each task, decoding performance dropped close to chance level (11.11%) when the classifiers were trained on perception trials and tested on WM trials or when the classifiers were trained on WM trials and tested on perception trials (Fig. 6a, b; SFig. 5a-d). In comparison, classifiers trained on one half of the WM trials and tested on the other half resulted in performance well above chance levels (Median Decoding Accuracy = 51.3%) (Fig. 6c; SFig. 5e, f). The latter indicates that our results were not an artifact of using different sets of trials for testing and training the classifiers, but were an effect of task type (perception vs WM).

We also conducted cross-epoch decoding for WM in which we trained and tested on combinations of cue, delay, and response epochs. Decoding performance was greatest when the classifiers were trained and tested with data from the same epoch and lowest when it was trained and tested on data from response and cue epochs (Train on cue - test on response: Median Decoding Accuracy = 11.0%; Train on response - test on cue: Median Decoding Accuracy = 12.3%) and when data was trained on the delay epoch and tested on the cue epoch (Median Decoding Accuracy = 17.0%) (Fig. 6d; SFig. 5g, h).

We also conducted cross-temporal decoding in which we trained and tested classifiers between congruent and incongruent time windows of 500 ms. These results indicate higher decoding accuracy when classifiers were trained and tested between temporally-near time windows within the same trial epoch. (SFig 5i). This data suggests that different neural activity profiles support LPFC neural codes for WM and perception.

## Discussion

By using complex virtual reality tasks, we were able to explore visuospatial WM and perception in naturalistic settings - incorporating natural eye movements and virtual navigation. We found that animals were able to accurately perform both tasks and identified distinct navigation strategies and eye movement behavior that occur during WM and perception. Whereas animals used a visually guided strategy in the perception task, they necessarily switched their strategy during WM. We also demonstrate the suitability of naturalistic WM tasks for neuronal recording in the LPFC, particularly those that allow for natural eye movements. We found that neurons in the primate LPFC are strongly tuned for target location during cue and delay epochs and that the amount of information during delay about target location remains consistent within the population of neurons on the single trial level. We also found that neuronal activity during fixation on target location is less predictive of target location during the delay epoch compared to the cue epoch indicating that eye position information does not contribute to decoding of target location during WM tasks. Information about target location encoded by the same neuronal population during the perception delay was not predictive of target location during the memory delay, indicating different patterns of population activity during perception and WM. Different population dynamics also exist between target encoding and memory epochs in the WM task.

### Influence of naturalistic task elements

One unique element of our task is the complex virtual environment in which it takes place since it contains non-relevant task stimuli. Based on the robust WM signals we describe, the LPFC may allow for the encoding of representations that are uniquely dissociated from distracting stimuli. Indeed, previous studies demonstrate that LPFC differs from areas such as the posterior parietal cortex where WM representations are perturbed by visual distractors (Suzuki, & Gottlieb, 2013; Jacob & Nieder, 2014). Evidence collected decades earlier from Malmo (1942) and Orbach and Fischer (1959) also report the importance of the PFC in maintaining WM representations in the presence of irrelevant incoming visual signals. However, we must be cautious when defining non-relevant stimuli, particularly in our virtual WM task where some of the elements of the environment (e.g., tree) may potentially be used as landmarks to estimate the target location during navigation.

Importantly, despite unconstrained eye movements, animals perform well on our WM task and the neuronal population maintains target selectivity and information about remembered location throughout the delay epoch. These findings may seem to contradict some previous literature showing that forced saccadic eye movements during memory delay reduces WM performance in human subjects (Postle, Idzikowski, Sala, Della, Logie, & Baddeley, 1999) and differentiates the LPFC from regions like the frontal eye fields where shifts in gaze disrupt WM signals (Balan, Ferrera, 2003). However, a distinction between our task and previous research is the production of forced versus naturally occurring saccades. Because the latter may be spontaneously and voluntarily triggered by the subjects, they may not interfere with performance in the same manner as task dependent saccades. Indeed, before the widespread use of the ODR and other oculomotor dependent tasks, simple delayed response tasks were used that displayed two targets and relied on an arm motor response through the use of the Wisconsin General Test Apparatus or button pressing. Although eye movements were not controlled in these classic experiments, studies reported neurons in the PFC that displayed clear delay activity and spatial selectivity (Fuster, & Alexander, 1971; Kojima, & Goldman-Rakic, 1982).

### Natural eye behavior and visuospatial working memory

Although the aim of our experimental paradigms was to approach natural behavior, potential concerns may arise surrounding the decision to not control eye position. For example, one may argue that animals would simply visually rehearse the target location by maintaining gaze fixation on the target of interest. We found substantial evidence against this behavioral strategy. Eye behavior differed between periods when the target was available compared to times when the target was unavailable like during the WM delay and response epochs. During WM delay, animals spent significantly less time looking onscreen, suggesting eye movement behavior that is less focused on specific elements in the environment such as target location. The number of fixations to target locations during WM delay only comprised 3% of fixations and there was no significant difference between the number of fixations on target between correct and incorrect trials, suggesting that fixation on target location during delay was not used as a successful behavioral strategy. From these results, one may infer the LPFC maintains an allocentric representation of the remembered location that is independent from gaze or fixation position. This issue, however, needs further exploration.

Using linear classifiers, we also identified that eye position on screen was significantly more predictive of target location during the cue epoch compared to the delay epoch. Classifiers that were trained on eye position data from the cue epoch and tested on eye position data from the delay epoch resulted in decoding accuracy below chance level suggesting different eye movement patterns between target encoding and memory maintenance. Moreover, in a recent study, we demonstrate that eye behavior remains unaffected by pharmaceutical manipulation that severely reduces WM coding and performance. In this study, despite significant changes to WM processing, gaze is equally as predictive of target location before and after systemic ketamine administration (Roussy et al, 2021b).

Saccade characteristics are influenced by external motivations like task reward (Takikawa, Kawagoe, Itoh, Nakahara, & Hikosaka, 2002). Increases in peak velocities have been observed for task-related saccades - when fixating on a target is needed for information processing - compared to saccades without a task related motivation (Bieg, Bresciani, Bülthoff, & Chuang, 2012). This increased saccadic speed may be used to gather task related information quicker. Saccades to target locations may be considered task relevant compared to non-target saccades, thus supporting correct task completion and reward. We found that saccades that land on targets versus those that land off target show a greater difference in velocity when the target is physically present during the cue epoch or perception task compared to when it is removed during the WM delay. In fact, there were no significant differences in saccade speed to targets compared to non-targets during the WM delay. This may suggest that saccades to target locations during memory delay were influenced less by task relevant motivation and information seeking than those made during the cue encoding period. Alternatively it may reflect the fact that visually guided saccades to a target show higher peak velocities than to an ‘empty’ location in space (Edelman, Valenzuela, Barton, 2006).

Another potential issue is contamination of WM signals by signals related to eye movement. We explored the amount of information contained by neural activity about target location during fixation on target locations during the cue and delay epochs. We found significantly lower decoding accuracy during the delay epoch compared to the cue epoch, suggesting that more information was available to the neuronal population when animals fixate on a target that is present compared to when the target is absent. Indeed, the decoding accuracy during the delay epoch was close to chance (25%), suggesting that animals did not receive substantial spatial information about target location during periods of target location fixation during delay. These results may be due to activation of visual neurons by the presence of a visual target during the cue epoch.

Although saccadic responses are seen in the PFC, the task and type of motor response required by the task has been shown to alter neuronal responses (Quintana, Yajeya, & Fuster. 1988; Sakagami, & Niki, 1994; Yajeya, Quintana, & Fuster, 1988; Johnston & Everling, 2006). Neuronal response to eye movements like saccades in the PFC are often identified during trials of tasks that are contingent on an oculomotor response. Neuronal responses to saccades are however notably absent when saccades are spontaneous and task independent such as during inter-trial intervals (Funahashi, 2014). Indeed, in a recent study, using the same virtual task, we analyzed the proportion of neurons that were tuned for saccade position in retinocentric and spatiocentric reference frames. Only 9% of neurons were tuned for retinocentric saccades and 11% for spatiocentric. More importantly, only 2% and 3% of neurons respectively were also tuned for remembered target location, providing further evidence for separate populations of neurons that code for eye position and remembered locations during WM tasks (Roussy et al.,2021b).

### Perception and working memory in area 8a and 9/46

The separation of perception and WM has been recognized since 1883 when neurological conditions were described in which patients exclusively lost either the ability to perceive objects or picture them in mind (Bernard, 1883; Behrmann, Moscovitch, & Winocur, 1994). Early lesion studies also point to a separation of these functions in LPFC in which large lesions consistently produced WM deficits while retaining perceptual discrimination functions (Reviewed in Roussy et al., 2021a). Moreover, pharmacological manipulations using muscimol and ketamine produce WM deficits without altering perceptual performance (Sawaguchi, & Iba, 2001; Roussy et al., 2021b).

Here, we found that population codes for perception and WM representations of target location are not interchangeable. This finding is supported by previous work from Mendoza-Halliday et al, who found separate populations of LPFC neurons that code for either perception or WM for visual motion direction (Mendoza-Halliday et al., 2017). After combining neurons into a pseudo-population, they further demonstrated that population activity patterns could decode whether neuronal representations were perceptual or mnemonic, suggesting different patterns of neuronal activity corresponding to each function. That study; however, used pseudo populations of neurons rather than simultaneously recorded neurons and did not use naturalistic virtual tasks. Our results expand on and validate the ones of that study.

How is it possible for the LPFC to represent perceived visual features without confounding WM representations? One possibility is that patterns of activity remain separate through the activation of perceptual, mnemonic, and mixed neurons. Activity patterns of perception and WM cells may help the brain monitor and discriminate between the internal (WM) and external (perception) representations. Abnormal patterns of activation may cause disruptions in internal and externally driven representations triggering hallucinations for example if perceptual neurons are activated without visual input. Interestingly, ketamine administration similarly disrupts patterns of activity during WM through disinhibition of neuron activity for non-preferred locations causing severe WM deficits (Roussy et al., 2021b).

### Conclusion

Our findings provide evidence of robust perceptual and WM representations in the macaque monkey LPFC during naturalistic tasks in virtual environments in which eye movements are unconstrained and the visual scene contains complex stimuli. We find minimal impact of natural eye movement on WM performance or neuronal coding for WM. Finally, we provide evidence for different neural codes for perceptual and mnemonic representations in the LPFC.

## Materials and Methods

The same two male rhesus macaques (Macaca mulatta) were used in both tasks (age: 10, 9; weight: 12, 10 kg).

### Ethics statement

Animal care and handling including basic care, animal training, surgical procedures, and experimental injections were pre-approved by the University of Western Ontario Animal Care Committee. This approval ensures that federal (Canadian Council on Animal Care), provincial (Ontario Animals in Research Act), and other national CALAM standards for the ethical use of animals are followed. Regular assessments for physical and psychological well-being of the animals were conducted by researchers, registered veterinary technicians, and veterinarians.

### Task

The current task takes place in a virtual environment that was created using Unreal Engine 3 development kit (UDK, May 2012 release; Epic Games). The nine targets were arranged in a 3 × 3 grid spaced approximately 0.5 seconds apart (movement speed during navigation was fixed). For the working memory task, the target is present only during the cue epoch. For the perception task, the target is present in the cue, delay, and response epochs. Detailed descriptions of this platform and the recording setup can be found in Doucet, Gulli, and Martinez-Trujillo, 2016.

### Experimental setup

During the task training period, animals were implanted with custom fit, PEEK cranial implants which housed the head posts and recording equipment (Neuronitek). See Blonde et al, 2018 for more information. Subjects performed all experiments while seated in a standard primate chair (Neuronitek) located in an isolated radiofrequency (RF) shielded room with the only illumination originating from the computer monitor. Animals were head posted during experiments and were delivered juice reward through an electronic reward integration system (Crist Instruments). The task was presented on a computer LDC monitor positioned 80 cm from the animals’ eyes (27” ASUS, VG278H monitor, 1024 × 768 pixel resolution, 75 Hz refresh rate, screen height equals 33.5 cm, screen width equals 45 cm). Eye position was tracked using a video-oculography system with sampling at 500 Hz (EyeLink 1000, SR Research).

### Microelectrode array implant

We chronically implanted two 10×10 microelectrode Utah arrays (96 channel, 1.5 mm in length, separated by at least 0.4 mm) (Blackrock Microsystems) in each animal located in the left LPFC (area 8a dorsal and ventral, anterior to the arcuate sulcus and on either side of the principal sulcus) (Petrides, 2005). Electrode arrays were placed and impacted approximately 1.5 mm into the cortex. Reference wires were placed beneath the dura and a grounding wire was attached between screws in contact with the pedestal and the border of the craniotomy.

### Processing of neuronal data

Neuronal data was recorded using a Cerebus Neuronal Signal Processor (Blackrock Microsystems) via a Cereport adapter. The neuronal signal was digitized (16 bit) at a sample rate of 30 kHz. Spike waveforms were detected online by thresholding at 3.4 standard deviations of the signal. The extracted spikes were semi-automatically resorted with techniques utilizing Plexon Offiine Sorter (Plexon Inc.). Sorting results were then manually refined. We collected behavioral data across 20 WM sessions (12 in NHP B, 8 in NHP T) and neuronal data from 19 WM sessions. Behavior was recorded from 19 perception sessions (14 in NHP B, 5 in NHP T). Neuronal data was analyzed from 13 sessions in which the WM and perception task were performed during the same session.

### Task performance

Percent of correct trials was calculated for both the WM and perception task. Response time was calculated for correct trials as the duration between the start of navigation and the time in which animals reach the correct target location. The task arena was divided into an 4 × 4 grid forming 16 area cells (Fig. 2d). The trajectory of the animal was calculated for each trial consisting of x and y coordinates sampled every 0.002 seconds. We calculated the number of samples that fell within each cell – this determined which cells the animals entered during navigation as well as how much of the total trajectory fell within each cell (related to time spent in cells). Our optimal trajectory measure is calculated by dividing the real length of the trajectory (the Euclidean distance from each x, y positional data point) by the true optimal distance (determined by the Euclidean distance from the start location to the target location for a particular trial). A value of 1 indicates the shortest possible (i.e., most optimal) trajectory length.

### Characterizing eye movement

The percent of eye data points on screen is calculated as the number of data points that fall within the screen limits divided by the total number of eye data points during a given epoch. Off screen data points occur when the animal looks outside of the defined screen limits or when the animal closes its eyes (i.e., during blinking).

We characterized eye movements as saccades, fixations, or smooth pursuits based on methods outlined in Corrigan et al. (2017). Eye movement data was first cleaned by removing blinks, periods of lost signal or corneal-loss spikes (occurs when corneal reflection is lost and regained). The clean eye signal was smoothed with a second-order Savitzky-Golay filter with a window of 11 samples. Saccades were identified by periods of high angular acceleration of the eye of at least 10 ms. Individual saccades were determined by an intersaccadic intervals of at least 40 ms. Saccade start and end points were determined by consistent direction and velocity considering a threshold of continuous change of > 20° for at least three samples, or an acute change of > 60° at one sample. Foveations were classified as fixations or smooth pursuits based on sample direction and ratios of distances. Dispersion of samples, consistently of direction, total path displacement, and the total spatial range were considered.

We calculated the percentage of total eye movement events classified as fixations or saccades for each epoch during WM and perception and the percentage of smooth pursuits for the response epoch.

### Main sequence calculation

The main sequence reflects the relationship between the amplitude of the saccade and the peak velocity of the eye rotation towards the saccade’s endpoint. Saccade amplitude and velocity can change based on the value of the saccade target (Bendiksby & Platt,2006) or alertness of the subject (Di Stasi, Catena, Cañas, Macknik, & Martinez-Conde,2013). To calculate the main sequence, we separated saccades into bins of 3° of amplitude, starting at 2° and computed the average peak velocity for each bin. Saccades within the same amplitude bins were matched between tasks to account for the influence of saccade start location and direction (direction with a tolerance of ±13°, and the starting location within 7°).

### Spatial tuning

Tuning for spatial location was computed in all units (3950, 3092 in NHP B, 858 in NHP T) in 19 WM sessions using Kruskal–Wallis one-way analysis of variance on epoch-averaged firing rates with target location as the independent variable. A neuron was defined as tuned if the test resulted in p < 0.05.

### Decoding target location from neuronal ensembles

We used a linear classifier (SVM) (Libsvm 3.14) (Fan, Chang, Hsieh, Wang, & Lin 2008) with 5-fold cross validation to decode target position from z-score normalized population-level responses using both single units and multiunits on a single trial basis. We grouped targets based on location in the virtual arena into three groups: right targets, center targets, and left targets leaving us with 3 classes (33.33% chance level). We used the best ensemble method detailed in Leavitt et al. (2017b), in which we determined the highest performing neuron, paired this neuron with all others in the population to achieve the best pair, and combined the best pair iteratively with all other neurons to form the best trio. This was repeated until we reached a best ensemble of 20 neurons or 16 neurons. The classifiers used firing rates calculated over 500 ms time windows. Decoding accuracy at each time window was compared to chance performance using t-tests.

### Gaze analysis

We calculated the total fixation time during the delay epoch as well as the fixation time on the trial specific target location for correct trials and incorrect trials. We compared the proportion of fixation time on the target location related to all fixation time during delay (target location fixation duration / total fixation duration) between correct and incorrect trials.

### Decoding target location using eye position

The screen was divided into 16 cells of equal dimensions. The number of foveations classified as fixations were calculated within each cell during the cue and delay epochs. We used a linear classifier (SVM) with 5-fold cross-validation to determine whether target location could be predicted by the number of fixations within each area of the screen under the assumption that animals gather information from the virtual environment during such fixation periods (Corrigan et al., 2017). This analysis was compared with a decoding analysis using neuronal ensembles utilizing the same number of features (16 neuron ensembles).

### Decoding eye position from neuronal data

We used a linear classifier (SVM) with 5-fold cross validation to decode eye position on screen based on neuronal firing rates during period of eye fixation. Four target locations were selected as part of this analysis since their locations were non-overlapping on screen. Fixation periods occurring in either the cue or delay epoch that fell within these regions were used. Short fixation periods were removed (amplitude < 6 ms). Firing rate was calculated for each neuron during each fixation period and were z-score normalized. Neuronal populations included single units and multiunits.

### Decoding target location for working memory and perception

We used a linear classifier (SVM) with 5-fold cross validation to decode target location (9 targets) based on population neuronal activity. We used 13 sessions in which animals performed both the WM and perception task so that we could use the same population of neurons. We altered training and testing conditions so that classifiers were either trained on population activity during congruent tasks or incongruent tasks (e.g., trained on WM and tested on perception).

We divided WM trials into two random and separate datasets and tested/ trained classifiers on one half of the trials and trained on the other half. For the WM task, we trained classifiers on either congruent or incongruent task epochs (e.g., train during cue and test during delay).

### Statistics

Additional statistical information is outlined in Table. S1.

## Author Contributions

M.R, J.M.T. L.P, R.A.G, B.C, and R.L designed the research. M.R, R.L, and A.J.S performed the research. M.R, B.C, and R.A.G contributed unpublished analytic tools. M.R analyzed data. M.R created figures. M.R and J.M.T wrote the paper.

## Data availability

Data supporting the findings of this study is available from the corresponding authors on reasonable request and will be fulfilled by M.R.

## Code availability

MATLAB codes used the analyze the data are available from M.R.

## Acknowledgments

We thank registered veterinary technicians Kim Thomaes and Rhonda Kersten from the University of Western Ontario for their assistance in surgery and animal care; Guillaume Doucet from the University of Ottawa for technical assistance related to Unreal Development Kit; Kevin Barker from Neuronitek for engineering equipment for our experiments; Jonathan C. Lau from the Division of Neurosurgery, University Hospital for providing advice regarding surgery and surgical planning; This work was supported by Canadian Institute of Health Research Project Grant; Natural Sciences and Engineering Research Council of Canada (NSERC); Ontario Graduate Scholarship; Chrysalis Foundation (London, Ontario). LP acknowledges salary support from the Tanna Schulich Endowment Chair for Neuroscience and Mental Health.

## Supplementary Figures

**Figure 1:**
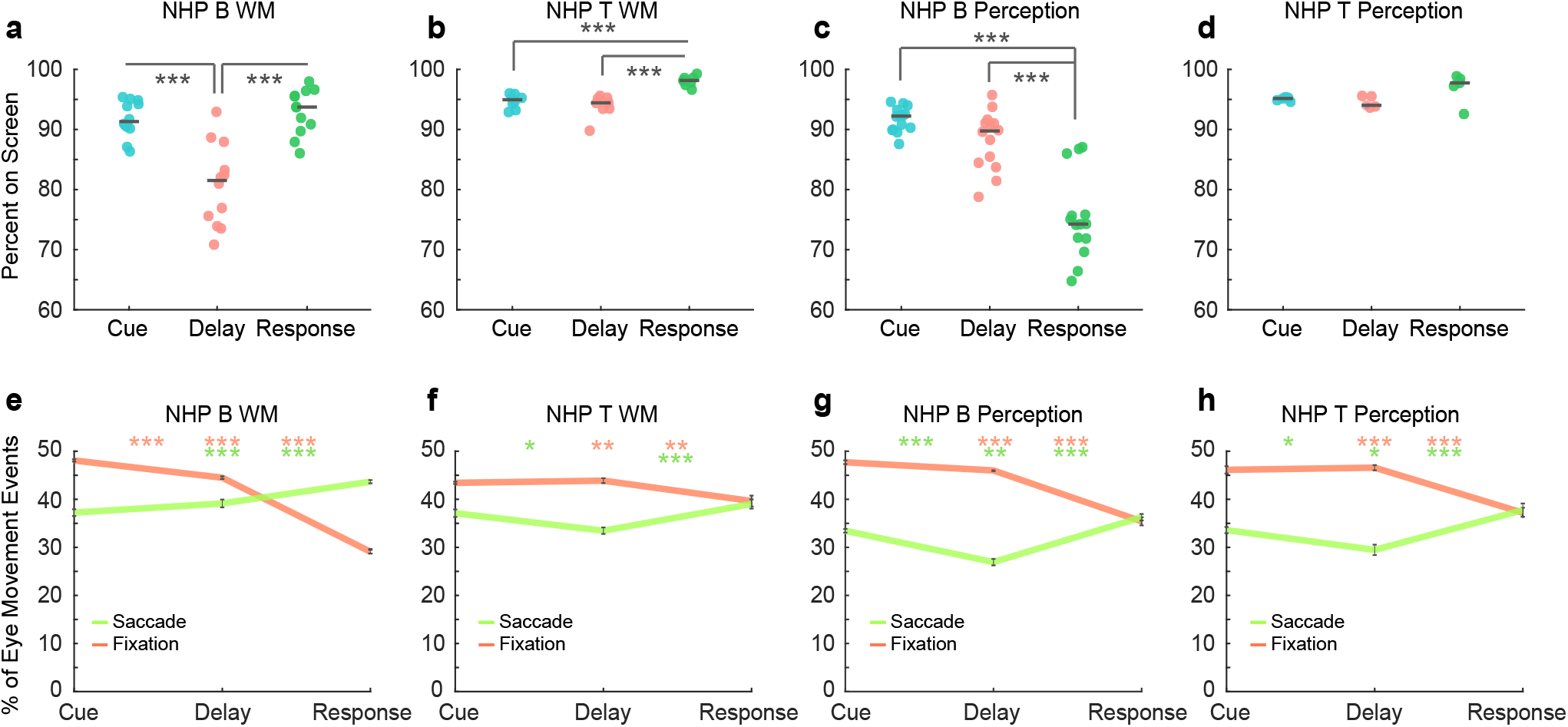
Eye Behavior for Each Animal. **a**. The percent of eye data points that fall within the boundaries of the screen during WM between trial epochs for NHP B. **b**. The percent of eye data points that fall within the boundaries of the screen during WM between trial epochs for NHP T. **c**. The percent of eye data points that fall within the boundaries of the screen during perception between trial epochs for NHP B. **d**. The percent of eye data points that fall within the boundaries of the screen during perception between trial epochs for NHP T. Dark grey lines represent mean values and each data point represents a session. **e**. Mean percent of eye movement events classified as fixations or saccades during WM between trial epochs for NHP B. **f**. Mean percent of eye movement events classified as fixations or saccades during WM between trial epochs for NHP T. **g**. Mean percent of eye movement events classified as fixations or saccades during perception between trial epochs for NHP B. **h**. Mean percent of eye movement events classified as fixations or saccades during perception between trial epochs for NHP T. Error bars are SEM. Asterisks on the left represent significance between the cue and delay epochs, asterisks in the middle represent significance between the cue and response epochs, and asterisks on the right represent significance between the delay and response epochs. Asterisk color corresponds to eye movement type. *p <* 0.01 = *, *p <* 0.001 = **, *p <* 0.0001 = ***.

**Figure 2:**
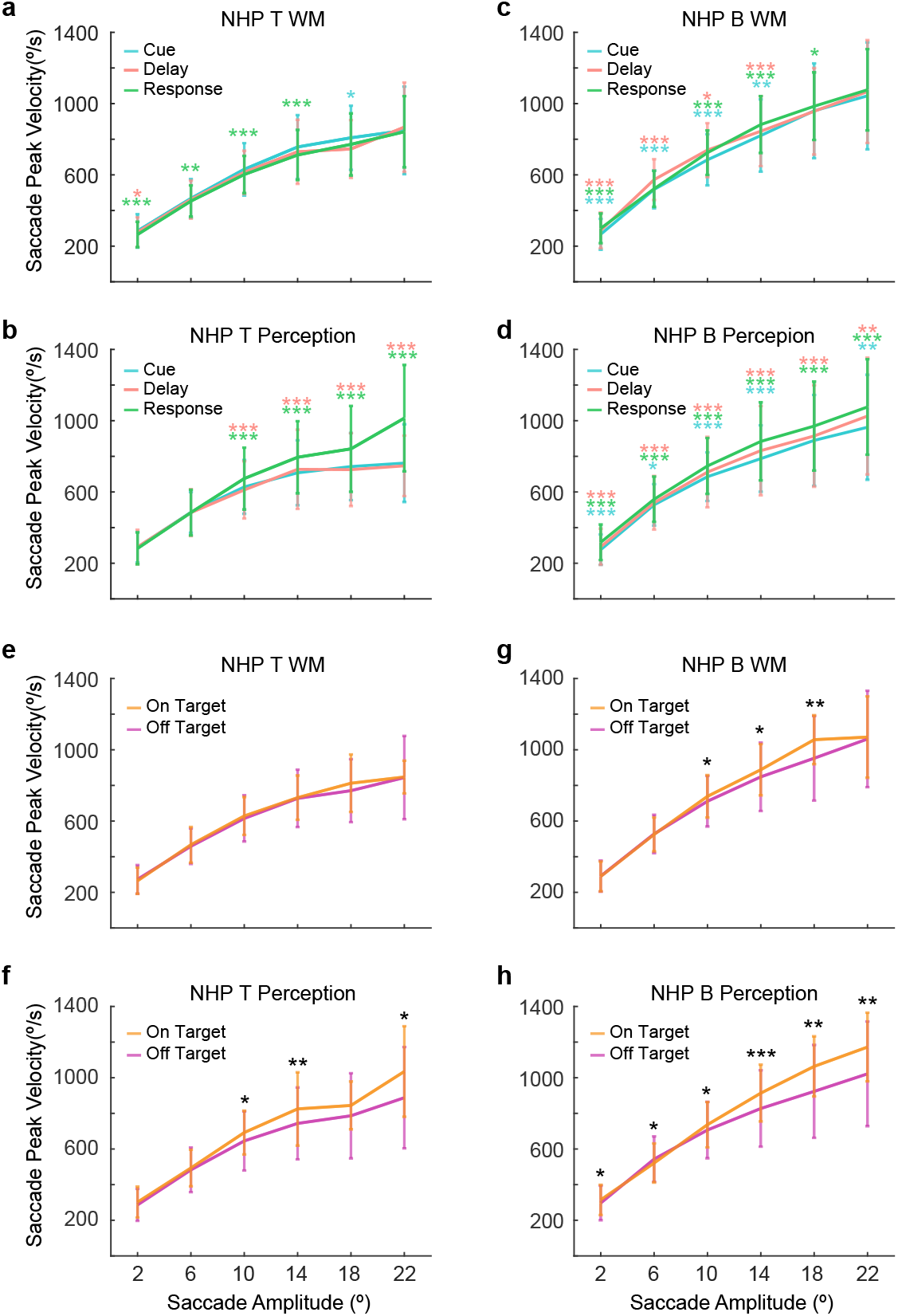
Saccade Behavior for Each Animal. **a**. Main sequence during WM between trial epochs for NHP T. **b**. Main sequence during perception between trial epochs for NHP T. **c**. Main sequence during WM between trial epochs for NHP B. **d**. Main sequence during perception between trial epochs for NHP B. Asterisks represent significance at each amplitude bin. Blue asterisks represent significance between the cue and delay epochs. Green asterisks represent significance between the cue and response epochs. Pink asterisks represent significance between the delay and response epochs. **e**. Main sequence during WM for saccades landing on and off targets for NHP T. **f**. Main sequence during perception for saccades landing on and off targets for NHP T. **g**. Main sequence during WM for saccades landing on and off targets for NHP B. **h**. Main sequence during perception for saccades landing on and off targets for NHP B. *p <* 0.01 = *, *p <* 0.001 = **, *p <* 0.0001 = ***.

**Figure 3:**
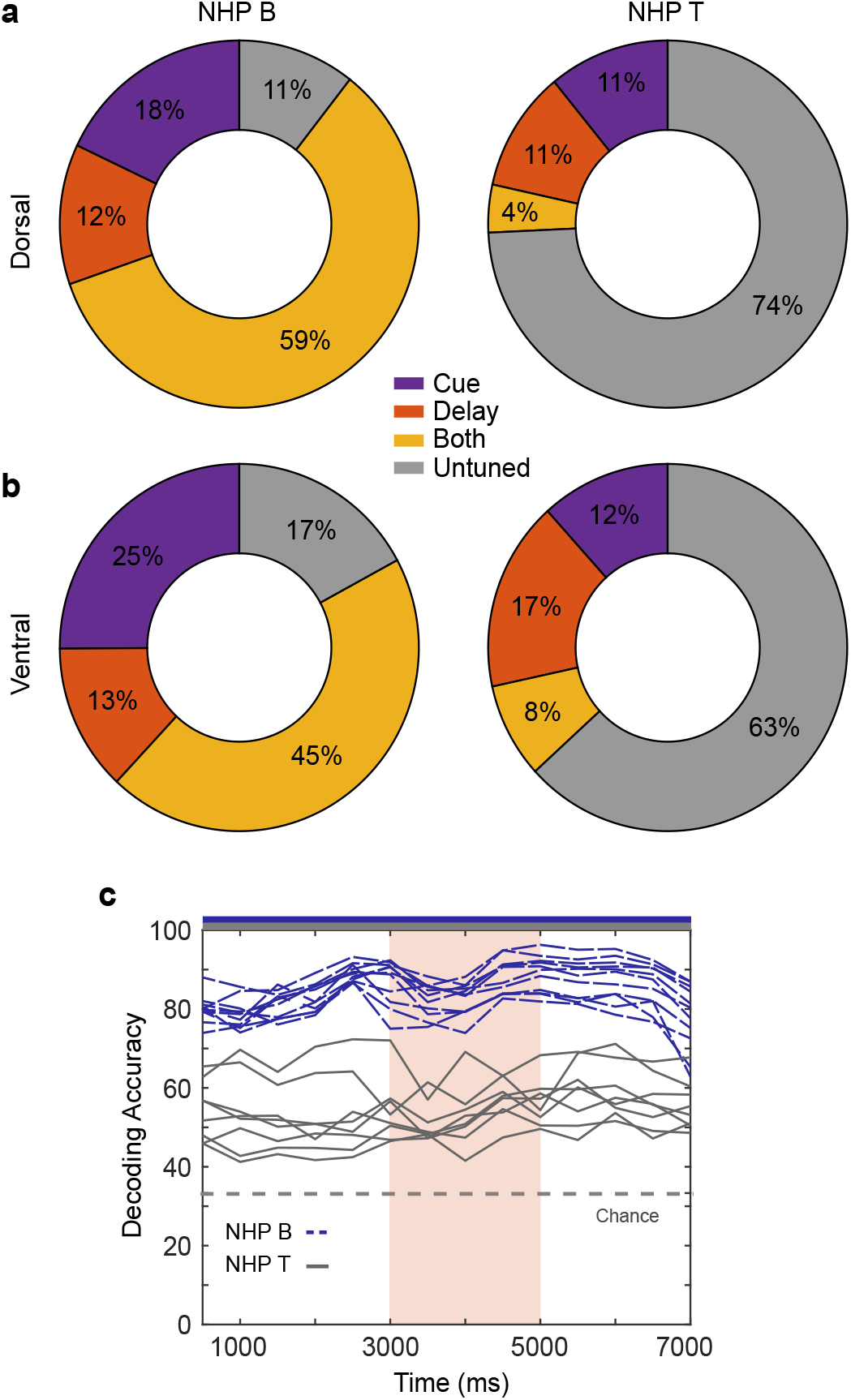
Neural Coding of Working Memory for Each Animal. **a**. Neural tuning during trial epochs for the dorsal array in NHP B and NHP T. **b**. Neural tuning during trial epochs for the ventral array in NHP B and NHP T. **c**. Decoding accuracy for neural decoding of target location. Each line represents data from one session. The orange column indicates the delay period. The grey dashed line indicates chance performance. Blue and grey bars on top of the figure represent significance from chance at each time window.

**Figure 4:**
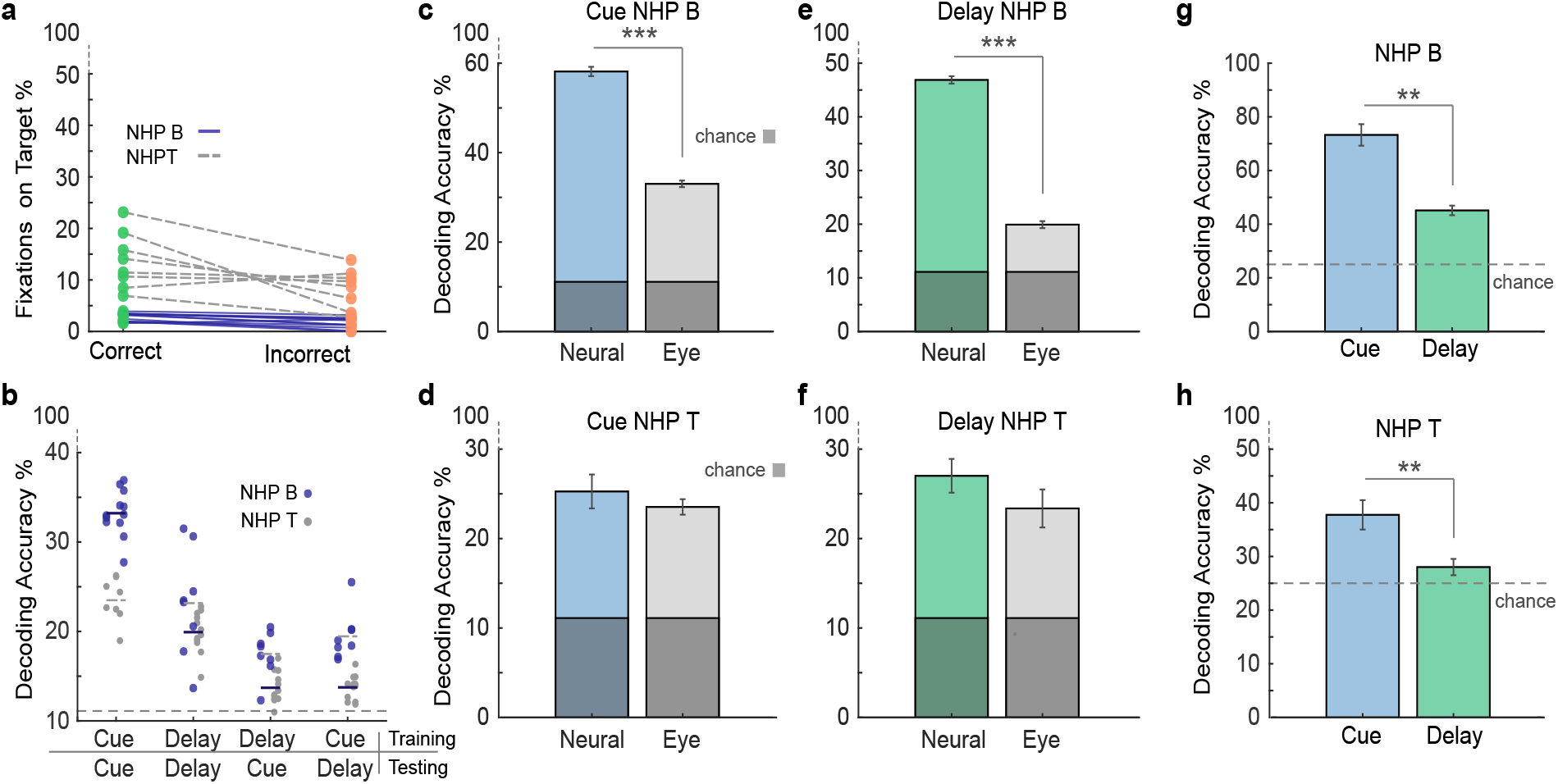
Eye Fixation Behavior for Each Animal. **a**. Percent of fixations on target for correct and incorrect trials. Dots represent data from individual sessions and lines connect correct and incorrect data from the same session. **b**. Decoding accuracy for predicting target location from the location of eye fixations on screen during the cue and delay epochs. Classifiers are trained on the first epoch listed in the label and tested on the second epoch. Dots represent data per session. **c**. Median decoding accuracy for target location based on neural data compared to eye position data for the cue epoch for NHP B. **d**. Median decoding accuracy for target location based on neural data compared to eye position data for the cue epoch for NHP T. **e**. Median decoding accuracy for target location based on neural data compared to eye position data for the delay epoch for NHP B. **f**. Median decoding accuracy for target location based on neural data compared to eye position data for the delay epoch for NHP T. **g**. Median decoding accuracy for eye position based on neural data during eye fixation periods for cue and delay epochs for NHP B. Grey dashed line represents chance performance. **h**. Median decoding accuracy for eye position based on neural data during eye fixation periods for cue and delay epochs for NHP T. Error bars are SEM. Grey dashed line represents chance performance. *p <* 0.01 = *, *p <* 0.001 = **, *p <* 0.0001 = ***.

**Figure 5:**
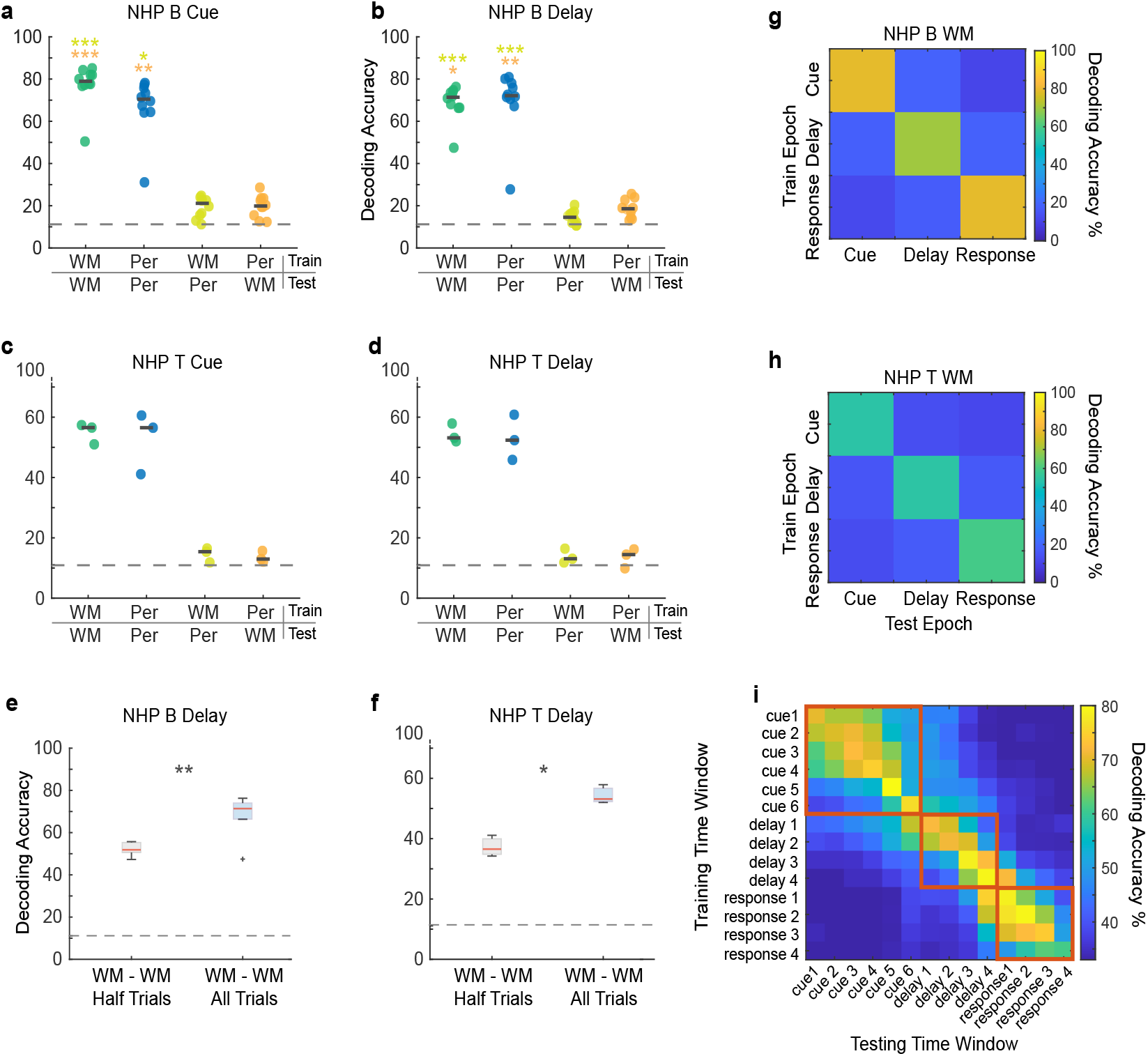
Cross-Trained Decoding Accuracy for Each Animal. **a**. Decoding accuracy for predicting target location for the perception and WM tasks during the cue epoch for NHP B. **b**. Decoding accuracy during the delay epoch for NHP B. **c**. Decoding accuracy during the cue epoch for NHP T. **d**. Decoding accuracy during the delay epoch for NHP T. Dark grey lines represent median values. The grey dashed line indicates chance performance. Classifiers are trained on the task that appears first in the x-axis label and tested on the task that appears second. **e**. Decoding accuracy with training and testing on one half of working memory trials during the delay epoch for NHP B. **f**. Decoding accuracy with training and testing on one half of working memory trials during the delay epoch for NHP T. The red lines represent median values and the bottom and top edges of the box indicate the 25th and 75th percentiles. The whiskers extend to non-outlier data points (within 1.5 std) and the outliers are plotted using ‘+’. **g**. Median decoding accuracy for WM when classifiers are cross-trained between epochs for NHP B. **h**. Median decoding accuracy for WM when classifiers are cross-trained between epochs for NHP T. **i**. Median decoding accuracy for WM when classifiers are cross-temporally crossed on 500 ms time windows throughout the trial period. Red rectangles surround classification results between time windows within epochs. *p <* 0.01 = *, *p <* 0.001 = **, *p <* 0.0001 = ***.

**Table 1.**
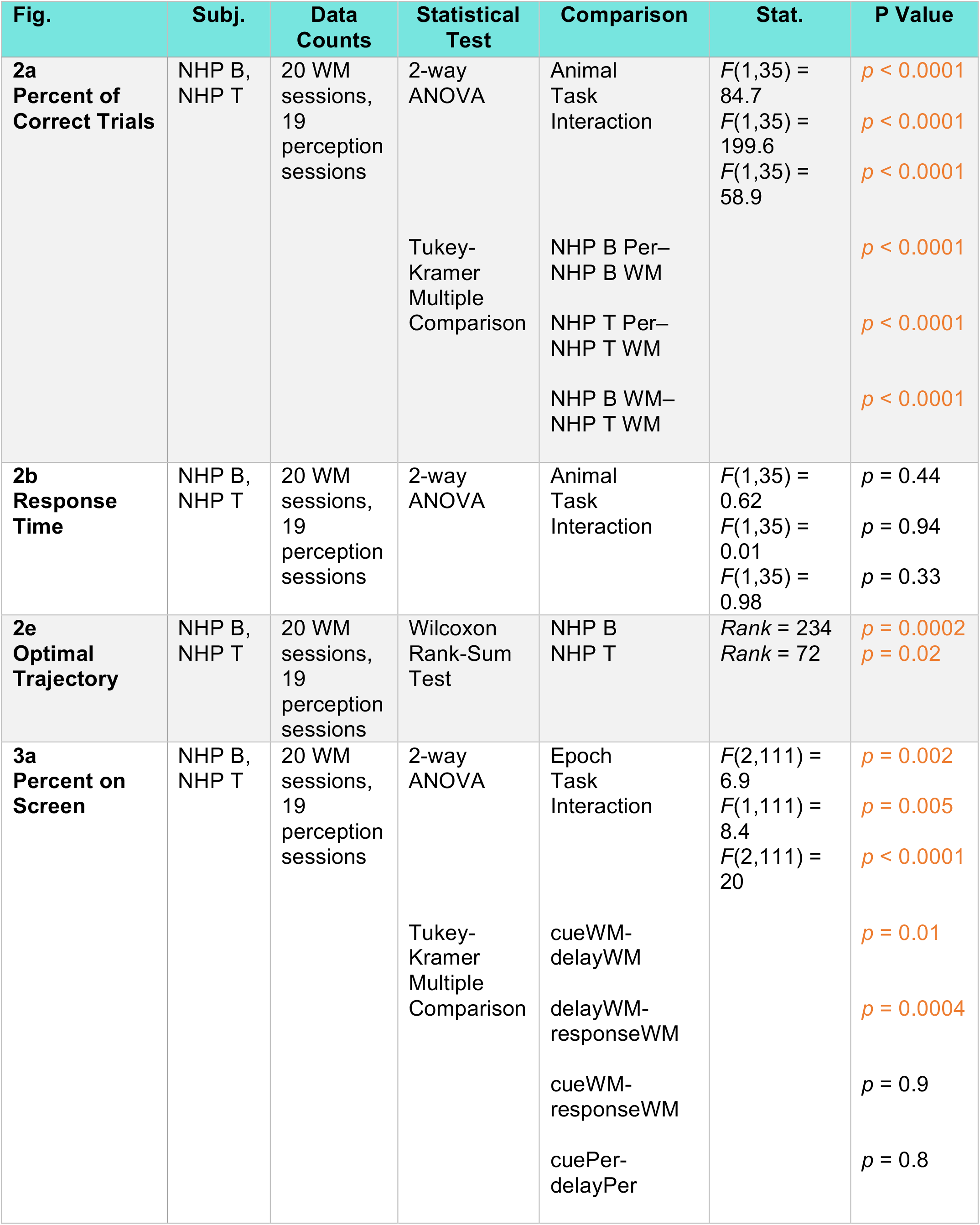
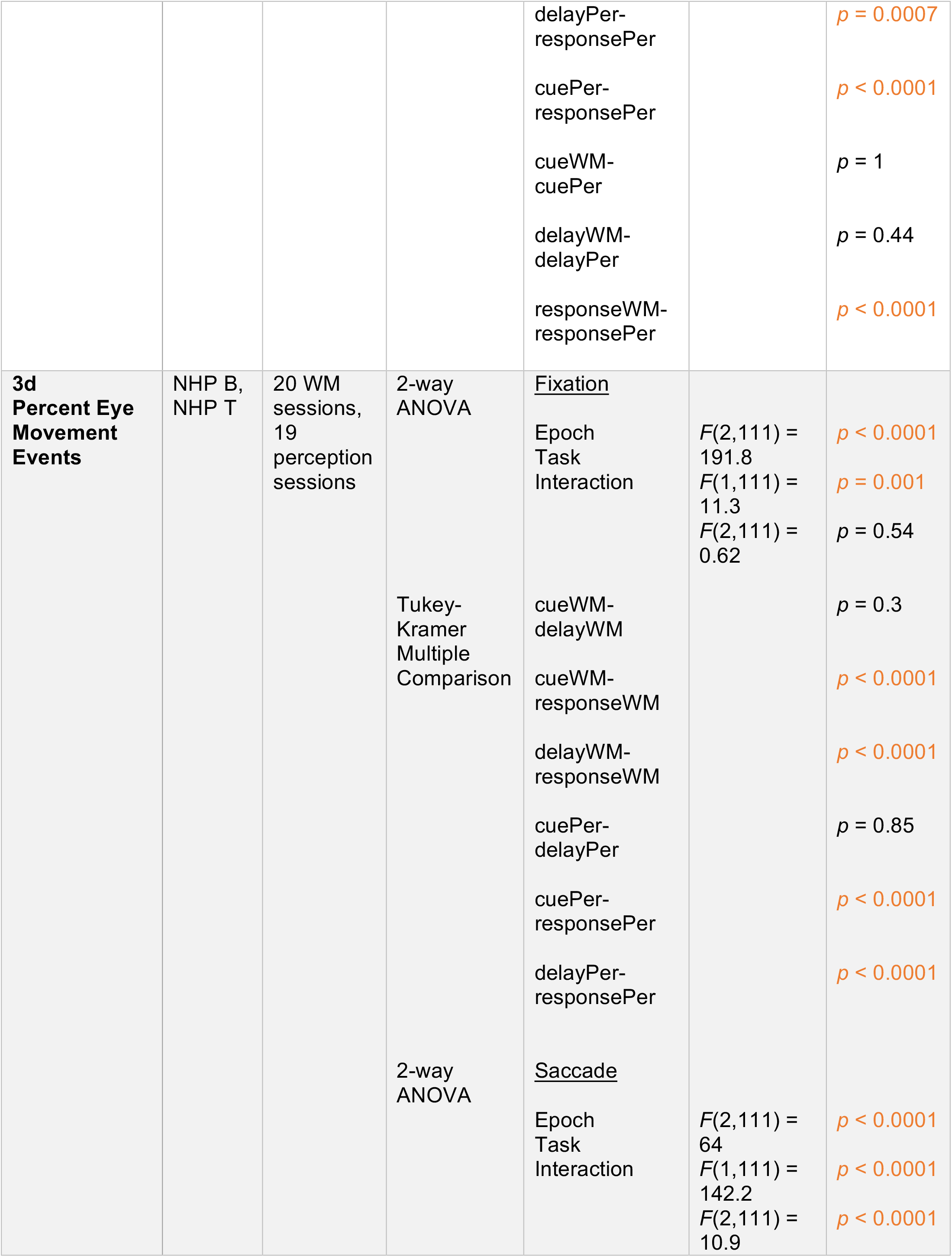
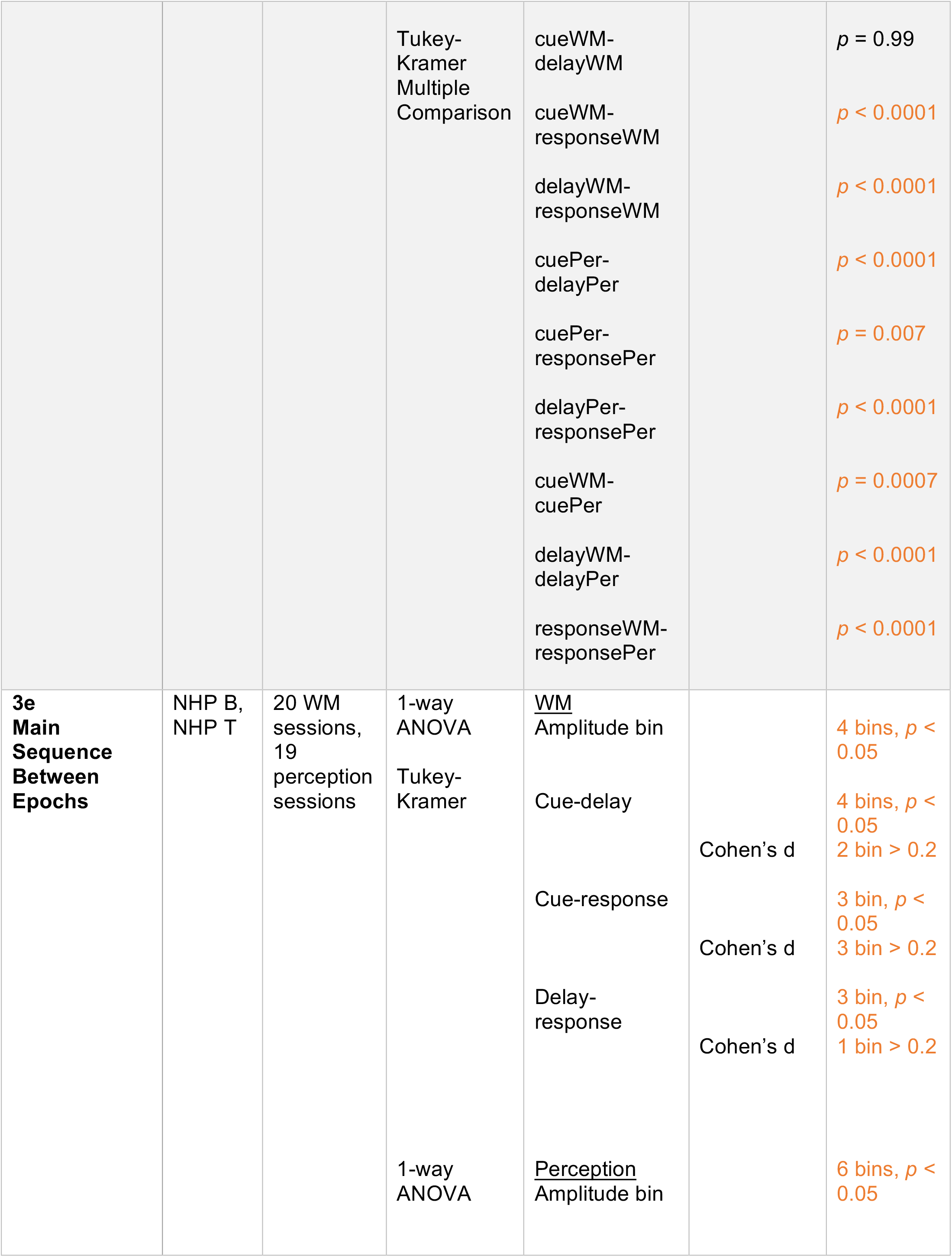
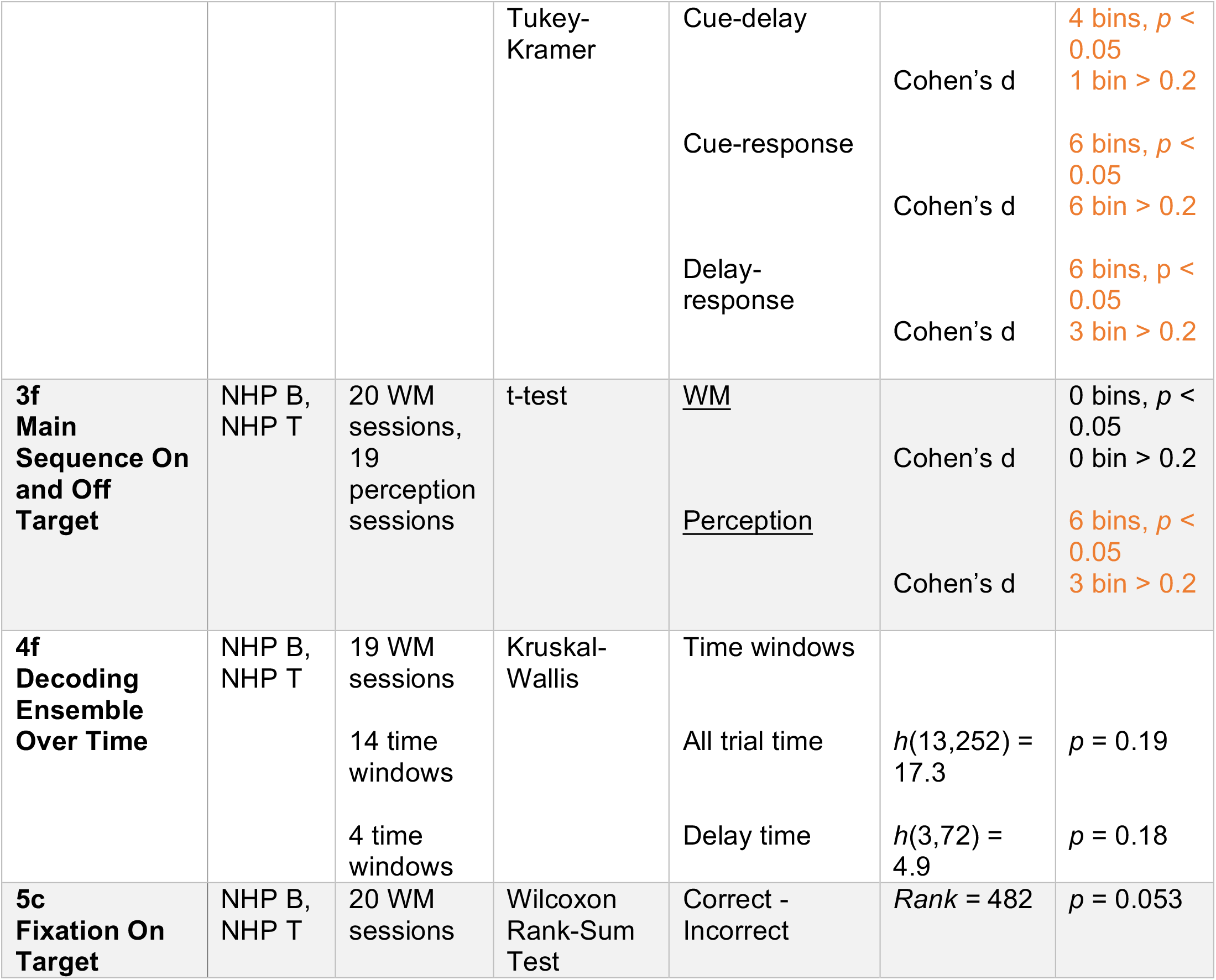
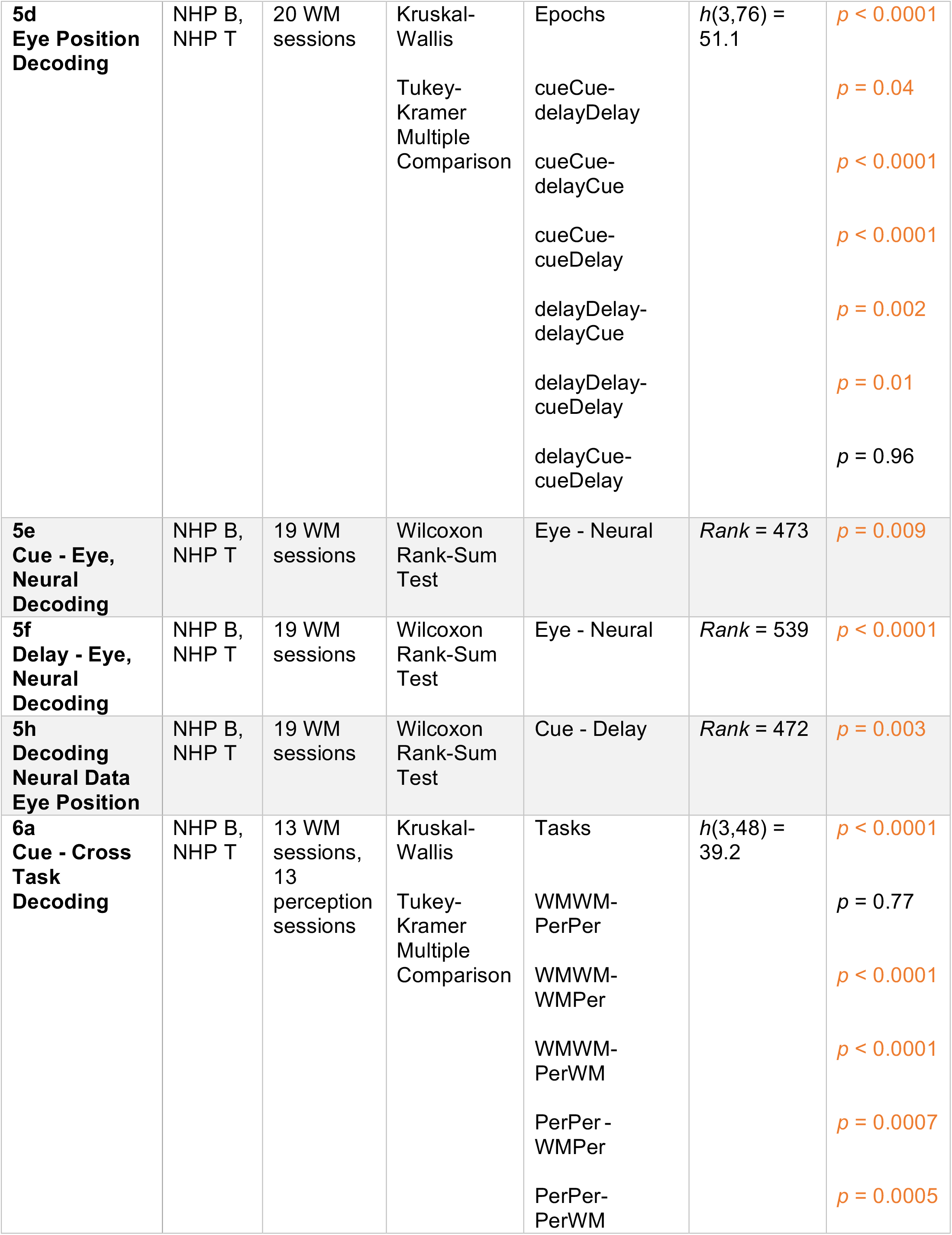
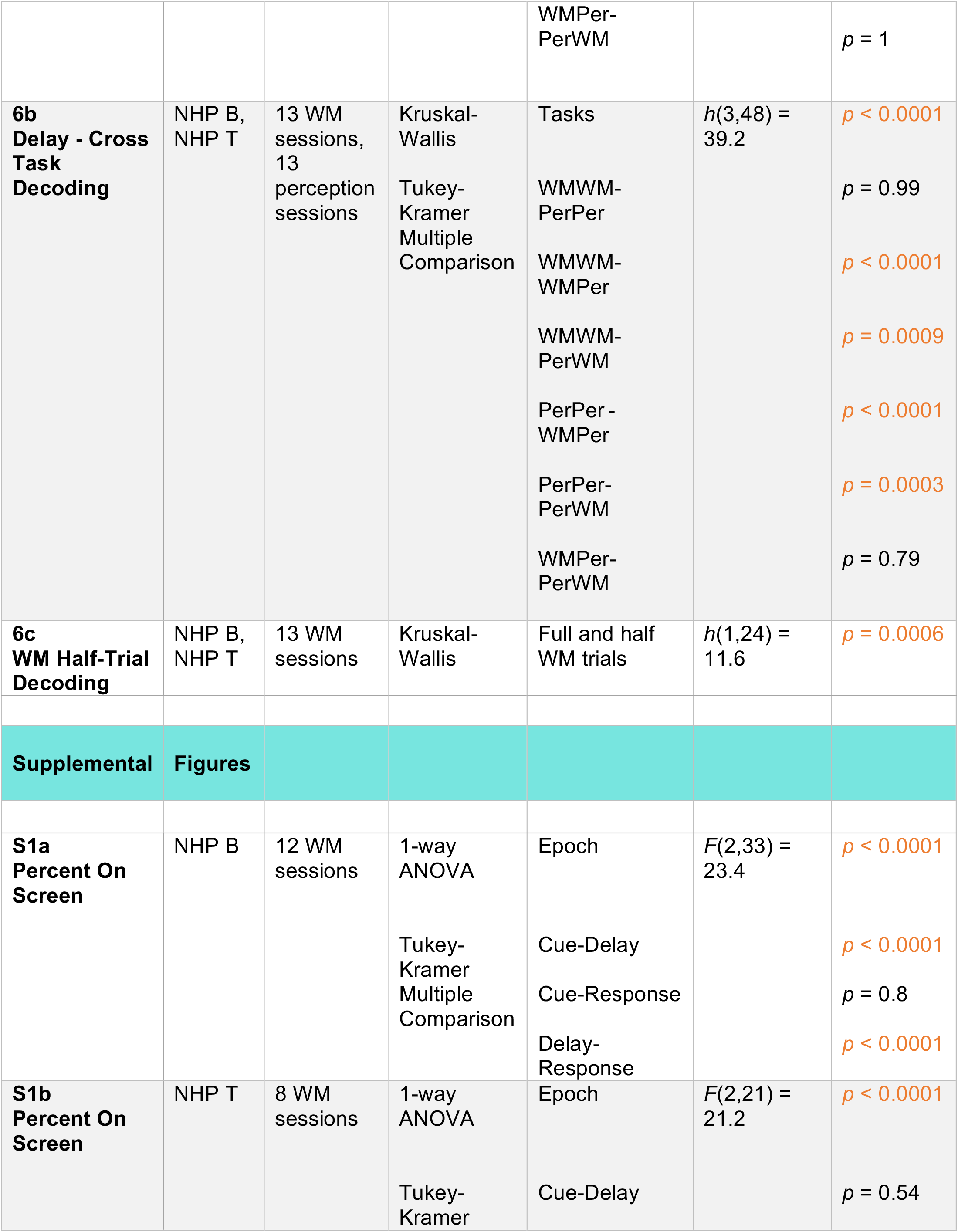
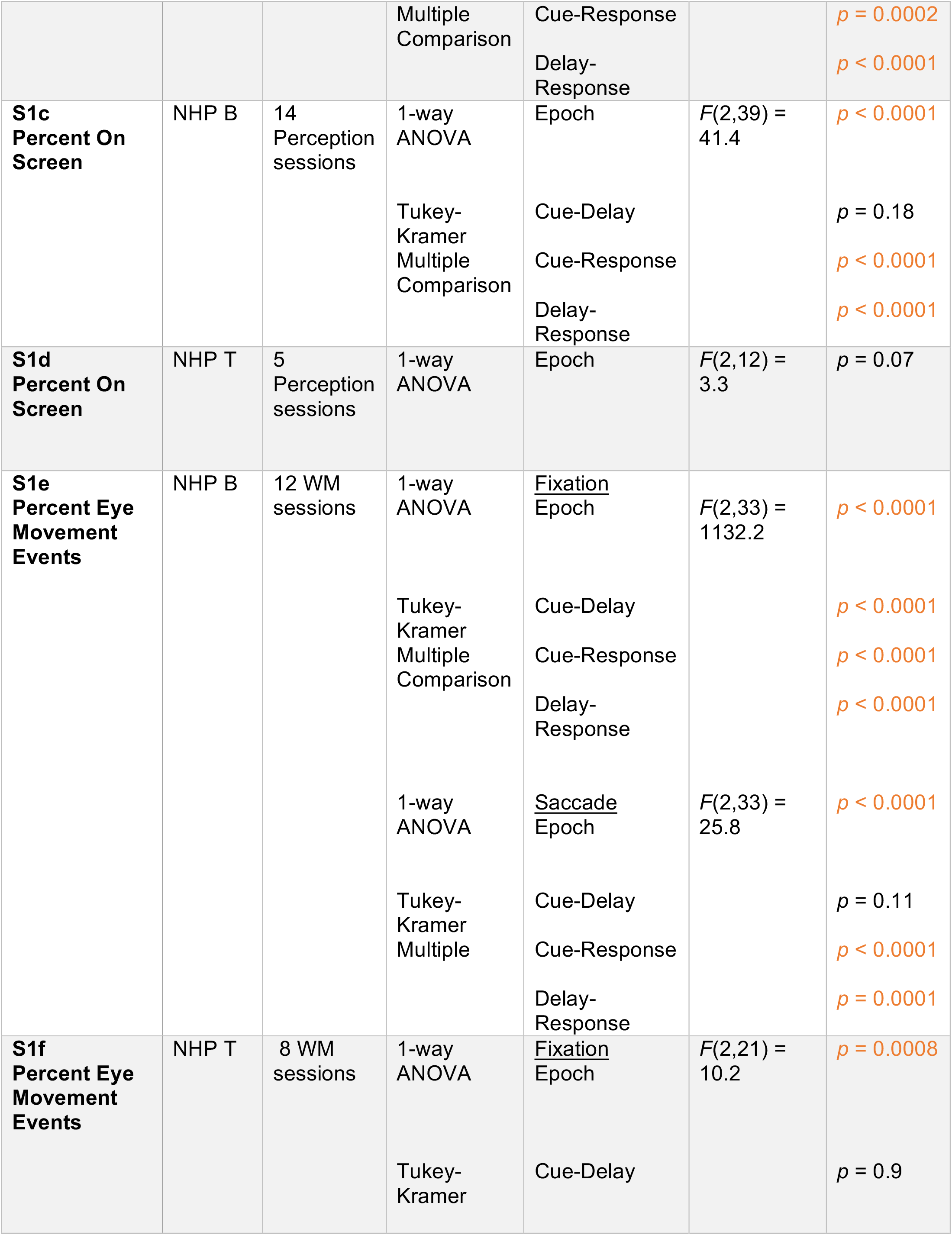
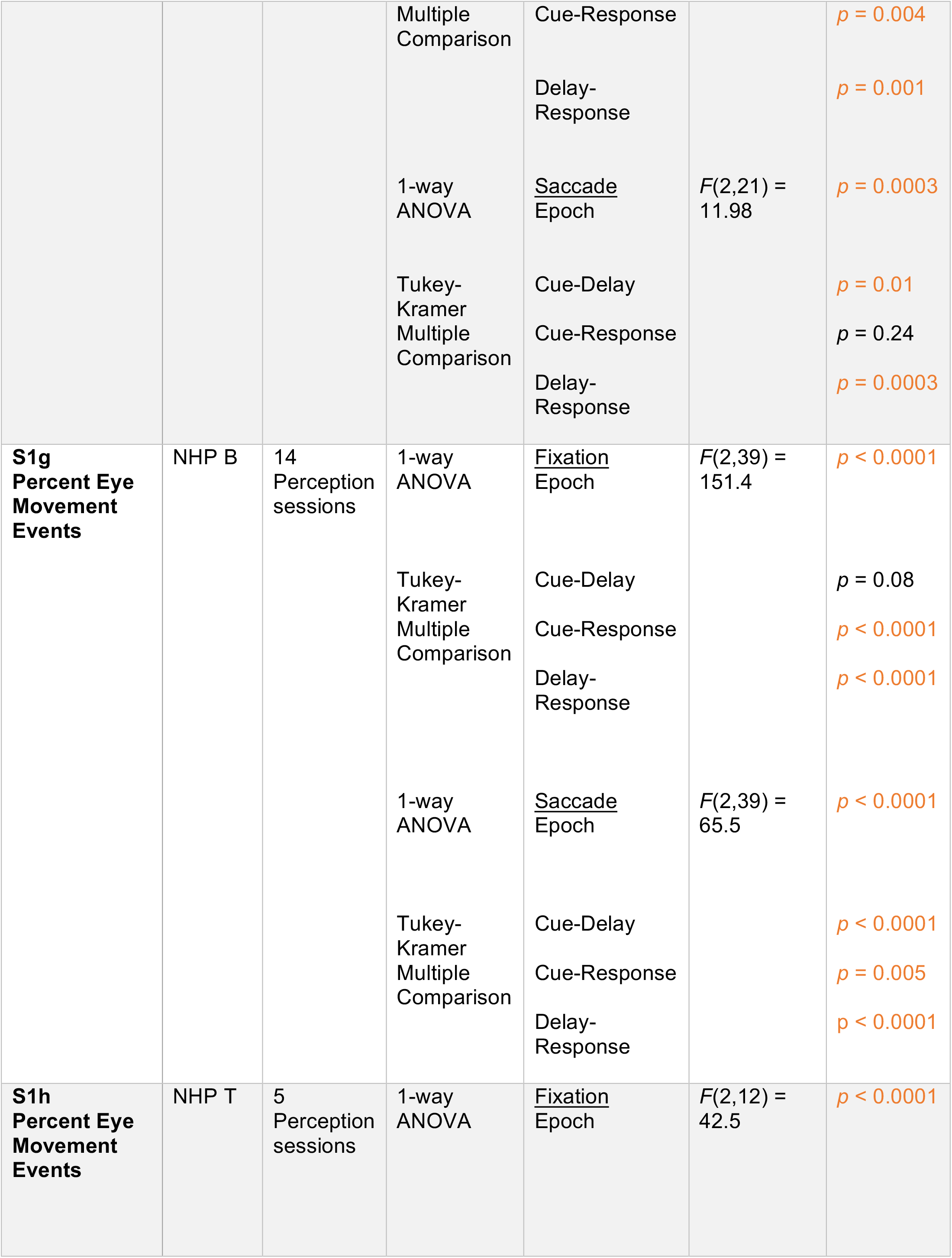
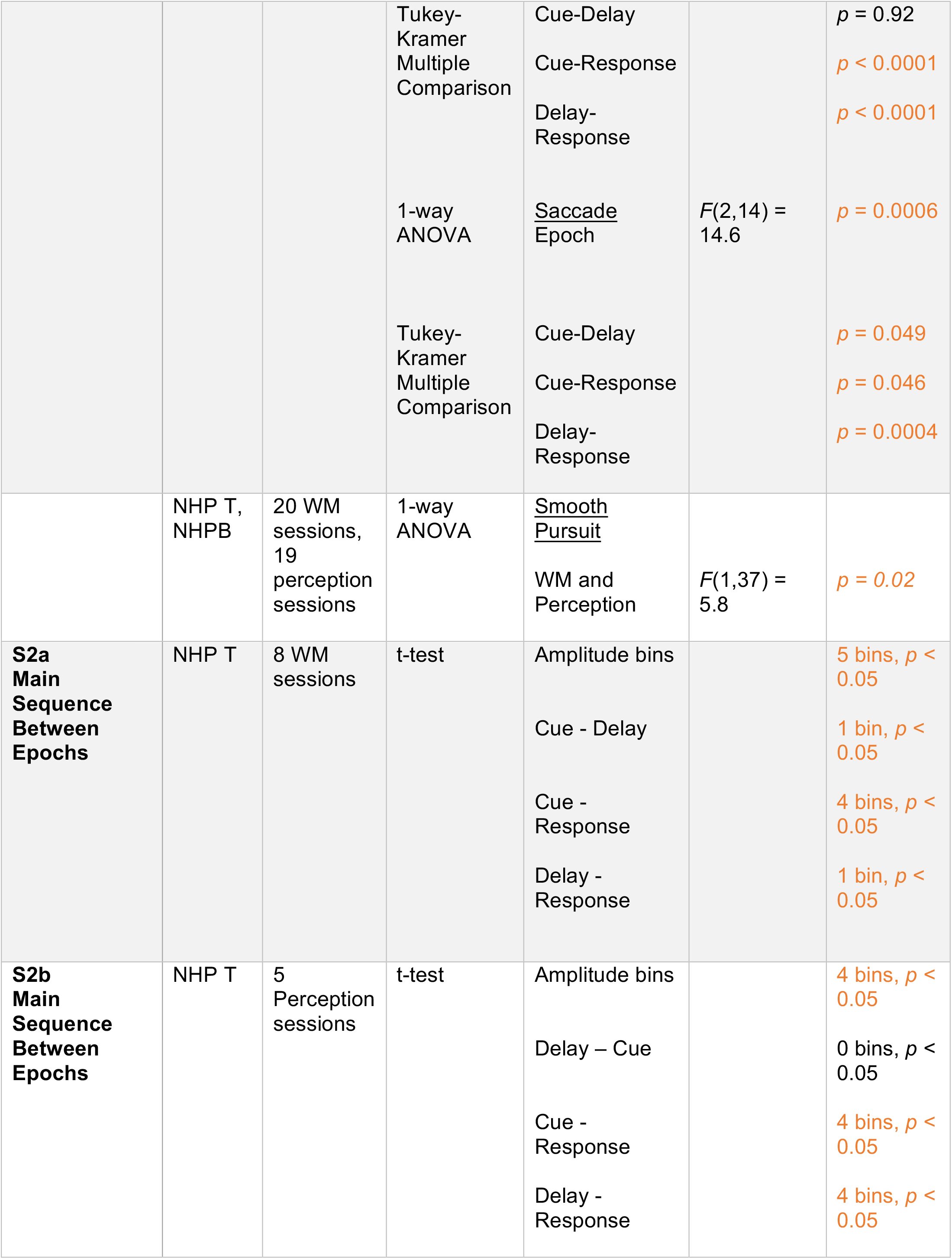
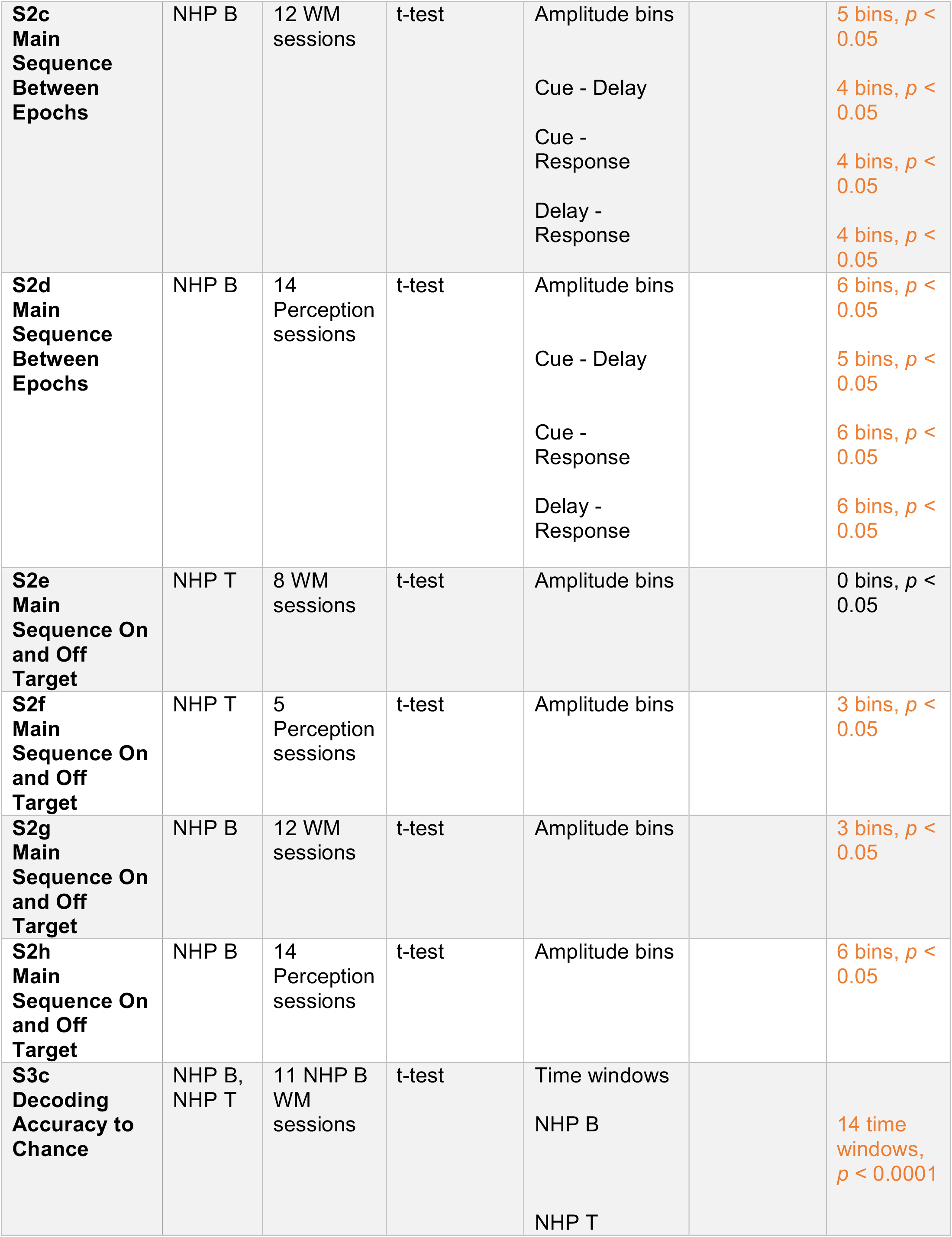
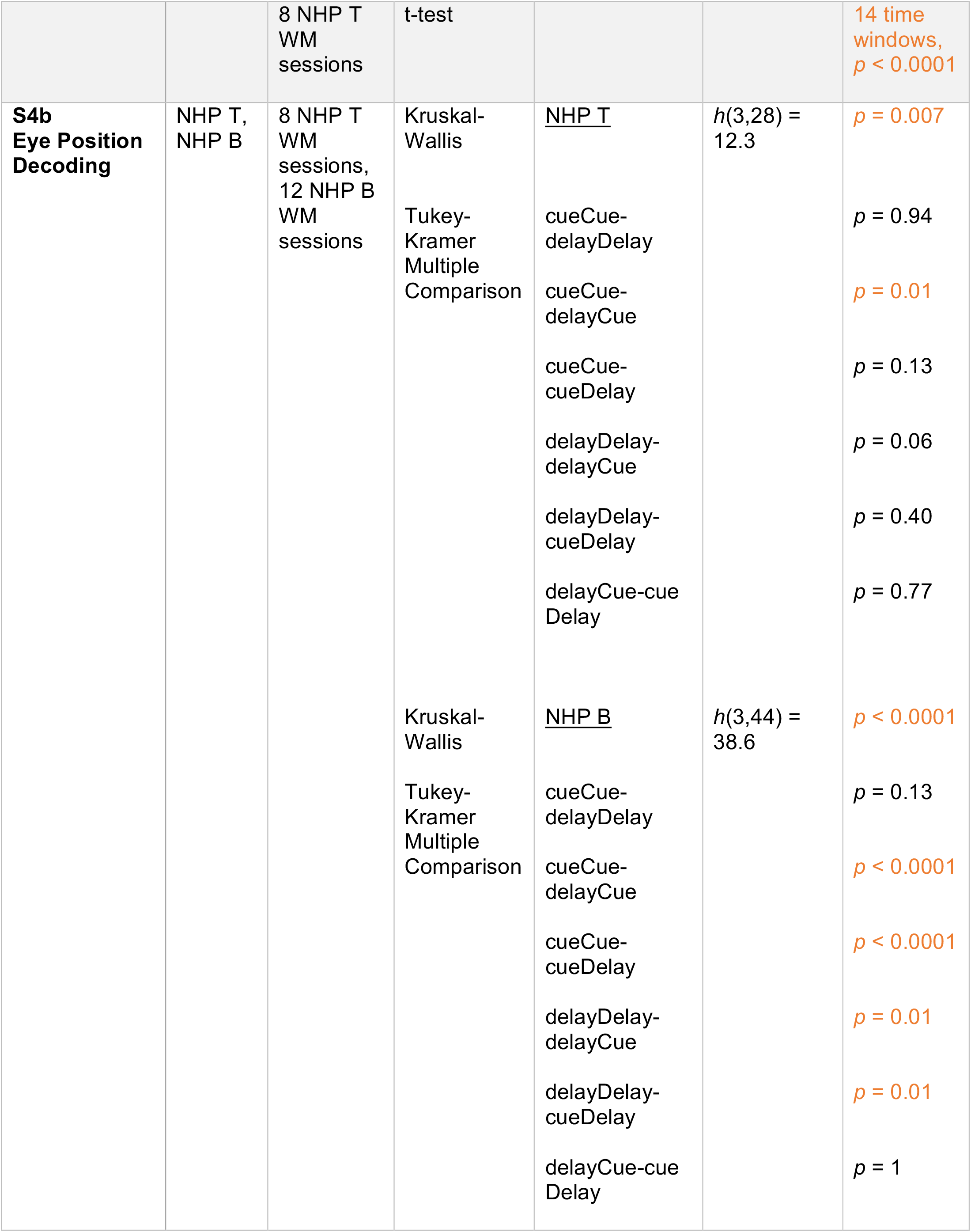
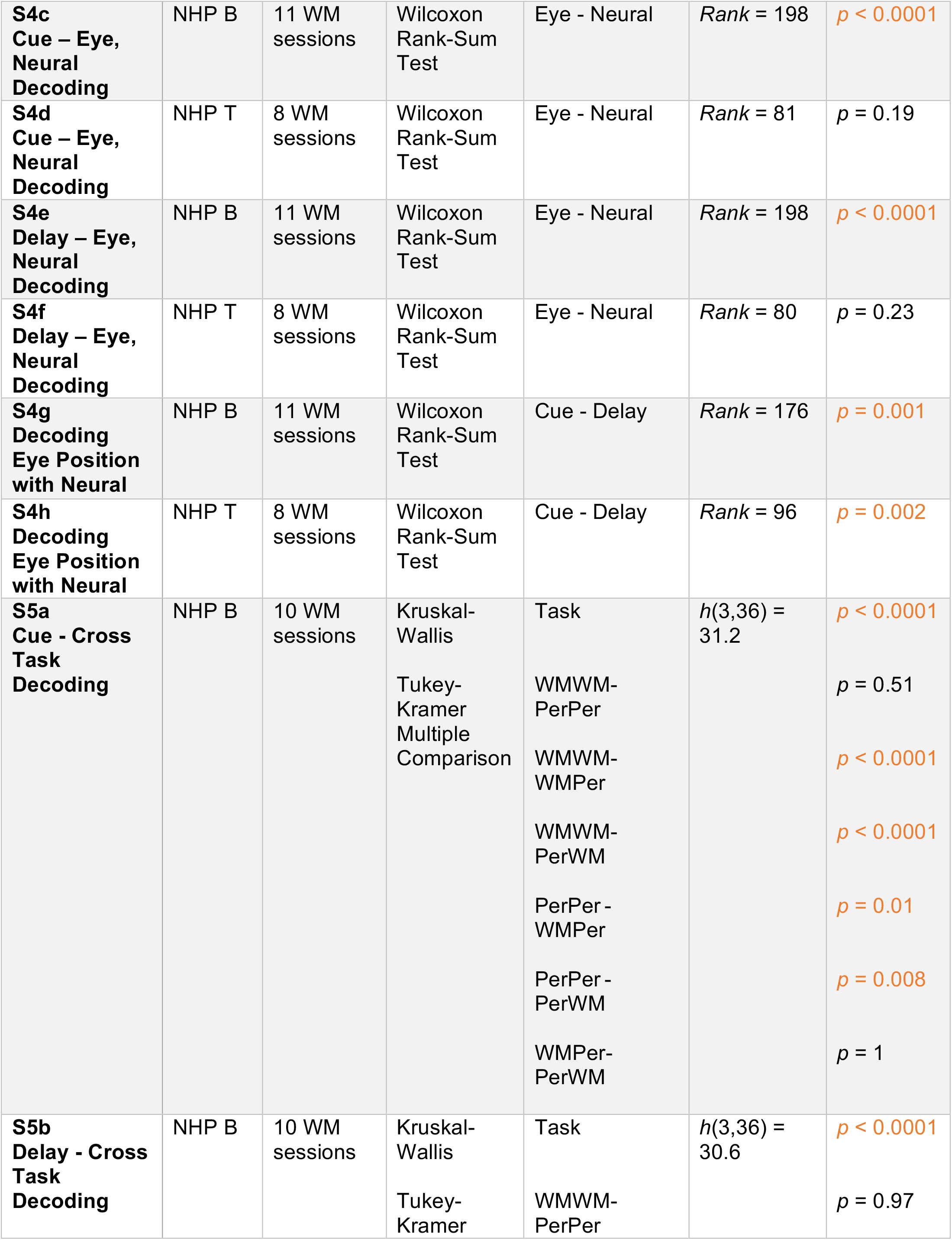
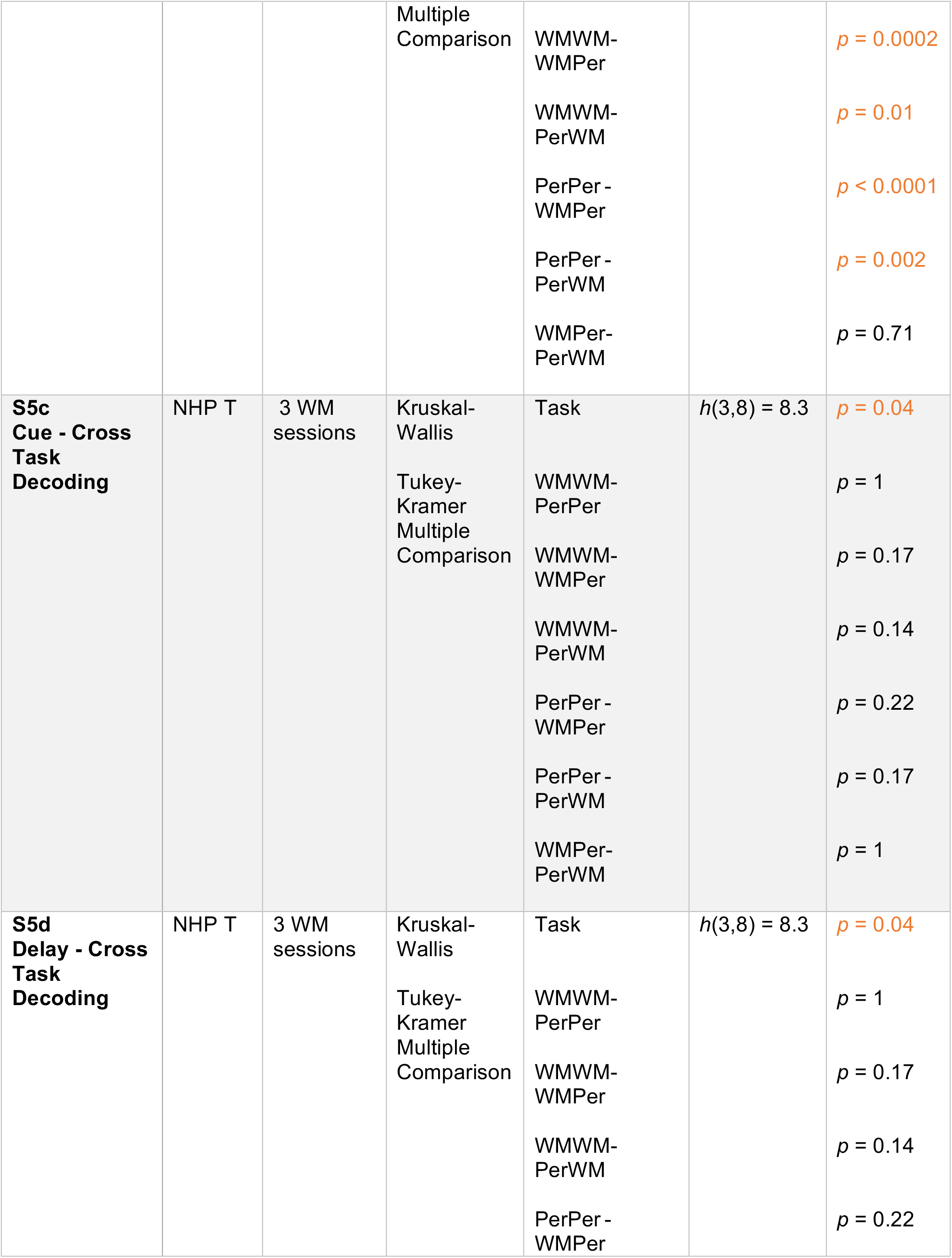
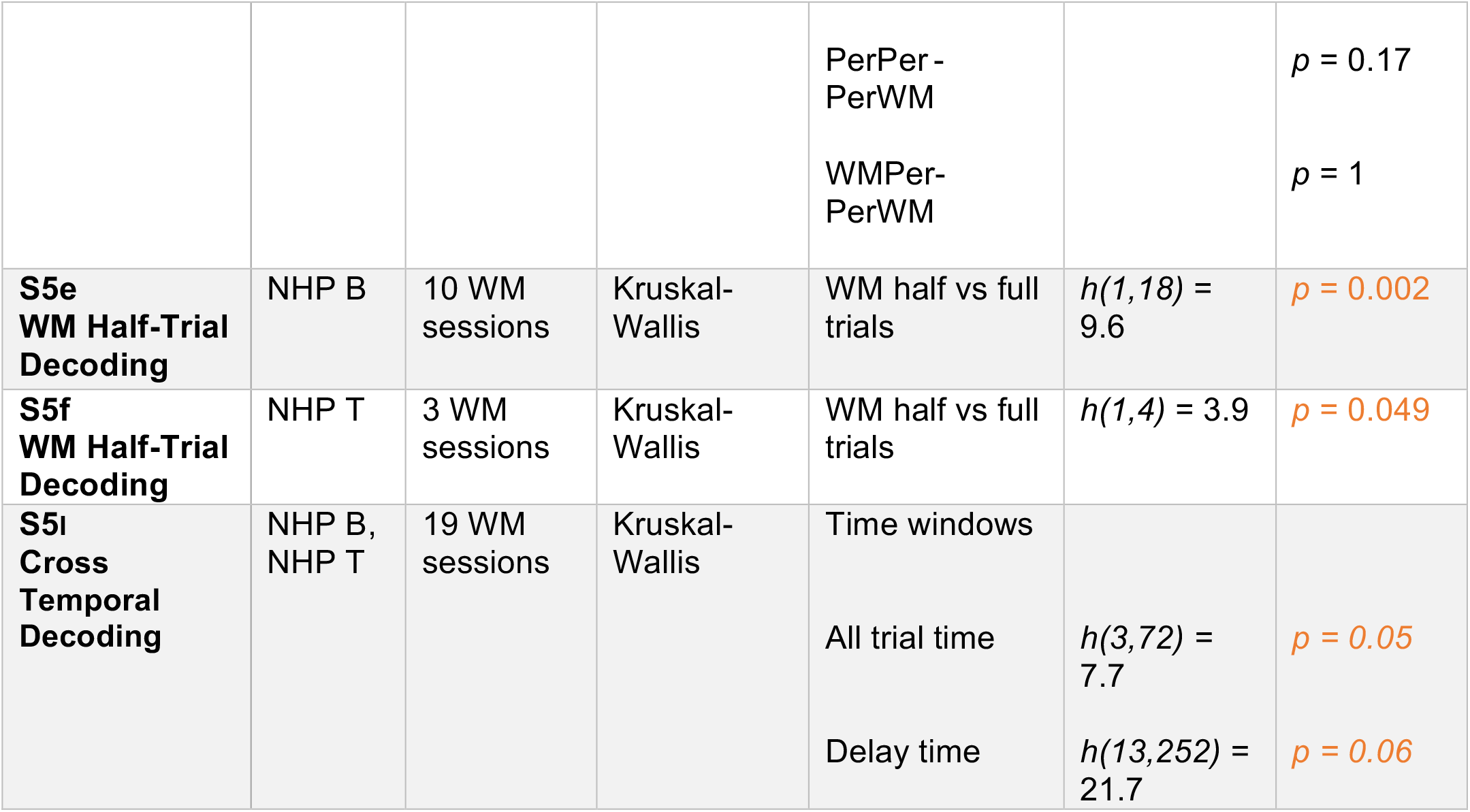
Statistics Reporting Table

## References

Baddeley, A. D. (1986). Working Memory. Clarendon Press/ Oxford University Press. Oxford, New York.

Balan, P. F., & Ferrera, V. P. (2003). Effects of gaze shifts on maintenance of spatial memory in macaque frontal eye field. The Journal of Neuroscience, 23 (13), 5446–5454. doi:10.1523/JNEUROSCI.23-13-05446.2003

Behrmann, M., Moscovitch, M., Winocur, G. (1994). Intact visual imagery and impaired visual perception in a patient with visual agnosia. Journal of Experimental Psychology: Human Perception and Performance, 20 (5), 1068–1087. doi:10.1037/0096-1523.20.5.1068

Bendiksby, M. S., & Platt, M. L. (2006). Neural correlates of reward and attention in macaque area LIP. Neuropsychologia, 44 (12), 2411–2420. doi:10.1016/J.NEUROPSYCHOLOGIA.2006.04.011

Bernard, D. A. F. (1883). Clinique des maladies nerveuses: un cas de suppression brusque et isolée de la vision mentale des signes et des objets: Formes et couleurs. Retrieved from https://gallica.bnf.fr/ark:/12148/bpt6k5463028m

Bieg, H. J., Bresciani, J. P., Bülthoff, H. H., & Chuang, L. L. (2012). Looking for discriminating is different from looking for looking’s sake. PLoS ONE, 7 (9). doi:10.1371/JOURNAL.PONE.0045445

Blonde, J. D., Roussy, M., Luna, R., Mahmoudian, B., Gulli, R. A., Barker, K. C., … Martinez-Trujillo, J. C. (2018). Customizable cap implants for neurophysiological experimentation. Journal of Neuroscience Methods, 304, 103–117. doi:10.1016/J.JNEUMETH.2018.04.016

Boulay, C. B., Pieper, F., Leavitt, M., Martinez-Trujillo, J., & Sachs, A. J. (2016). Single-trial decoding of intended eye movement goals from lateral prefrontal cortex neural ensembles. Journal of Neurophysiology, 115, 486–499. doi:10.1152/jn.00788.2015

Bullock, K. R., Pieper, F., Sachs, A. J., & Martinez-Trujillo, J. C. (2017). Visual and presaccadic activity in area 8Ar of the macaque monkey lateral prefrontal cortex. Journal of Neurophysiology, 118 (1), 15–28. doi:10.1152/JN.00278.2016

Constantinidis, C., Funahashi, S., Lee, D., Murray, J. D., Qi, X. L., Wang, M., & Arnsten, A. F. T. (2018). Persistent spiking activity underlies working memory. The Journal of Neuroscience, 38 (32), 7020–7028. doi:10.1523/JNEUROSCI.2486-17.2018

Corrigan, B. W., Gulli, R. A., Doucet, G., & Martinez-Trujillo, J. C. (2017). Characterizing eye movement behaviors and kinematics of non-human primates during virtual navigation tasks. Journal of Vision, 17 (12). doi:10.1167/17.12.15

Di Stasi, L. L., Mccamy, M. B., Catena, A., Macknik, S. L., Cañas, J. J., & Martinez-Conde, S. (2013). Microsaccade and drift dynamics reflect mental fatigue. European Journal of Neuroscience, 38 (3), 2389–2398. doi:10.1111/EJN.12248

Doucet, G., Gulli, R. A., & Martinez-Trujillo, J. C. (2016). Cross-species 3D virtual reality toolbox for visual and cognitive experiments. Journal of Neuroscience Methods, 266, 84–93. doi:10.1016/J.JNEUMETH.2016.03.009

Edelman, J. A., Valenzuela, N., & Barton, J. J. (2006). Antisaccade velocity, but not latency, results from a lack of saccade visual guidance. Vision Research, 46 (8-9), 1411–1421. doi:10.1016/j.visres.2005.09.013

Fan, R.-E., Chang, K.-W., Hsieh, C.-J., Wang, X.-R., & Lin, C.-J. (2008). LIBLINEAR: A Library for Large Linear Classification. Journal of Machine Learning Research, 9, 1871–1874. doi:10.1145/1390681.1442794

Funahashi, S., Bruce, C. J., & Goldman-Rakic, P. S. (1989). Mnemonic coding of visual space in the monkey’s dorsolateral prefrontal cortex. Journal of Neurophysiology, 61 (2), 331–349. doi:10.1152/JN.1989.61.2.331

Funahashi, S. (2014). Saccade-related activity in the prefrontal cortex: Its role in eye movement control and cognitive functions. Frontiers in Integrative Neuroscience, 8. doi:10.3389/FNINT.2014.00054

Fuster, J. M., & Alexander, G. E. (1971). Neuron activity related to short-term memory. Science, 173 (3997), 652–654. doi:10.1126/SCIENCE.173.3997.652

Goldman-Rakic, P. S. (1994). Cellular basis of working memory. Neuron,14 (3), 477–485, doi:10.1016/0896-6273(95)90304-6.

Hasegawa, R., Sawaguchi, T., Kubota, K., & Fuster, K. (1998). Monkey prefrontal neuronal activity coding the forthcoming saccade in an oculomotor delayed matching-to-sample task. Journal of Neurophysiology, 79 (1), 322–333.. doi:10.1152/jn.1998.79.1.322

Jacob, S. N., & Nieder, A. (2014). Complementary roles for primate frontal and parietal cortex in guarding working memory from distractor stimuli. Neuron, 83 (1), 226–237. doi:10.1016/j.neuron.2014.05.009

Johnston, K., & Everling, S. (2006). Neural activity in monkey prefrontal cortex is modulated by task context and behavioral instruction during delayed-match-to-sample and conditional prosaccade-antisaccade tasks. Journal of Cognitive Neuroscience, 18 (5), 749–765. doi:10.1162/jocn.2006.18.5.749

Kojima, S., & Goldman-Rakic, P. S. (1982). Delay-related activity of prefrontal neurons in rhesus monkeys performing delayed response. Brain Research, 248 (1), 43–50. doi:10.1016/0006-8993(82)91145-3

Leavitt, M. L., Mendoza-Halliday, D., & Martinez-Trujillo, J. C. (2017a). Sustained activity encoding working memories: Not fully distributed. Trends in Neurosciences, 40 (6), 328–346. doi:10.1016/J.TINS.2017.04.004

Leavitt, M. L., Pieper, F., Sachs, A. J., & Martinez-Trujillo, J. C. (2017b). Correlated variability modifies working memory fidelity in primate prefrontal neuronal ensembles. PNAS, 114 (12), 2494–2503. doi:10.1073/pnas.1619949114

Leavitt, M. L., Pieper, F., Sachs, A. J., & Martinez-Trujillo, J. C. (2018). A quadrantic bias in prefrontal representation of visual-mnemonic space. Cerebral Cortex, 28 (7), 2405–2421. doi:10.1093/cercor/bhx142

Malmo, R. B. (1942). Interference factors in delayed response in monkeys after removal of frontal lobes. Journal of Neurophysiology, 5, 295–308. doi:10.1152/jn.1942.5.4.295

Mendoza-Halliday, D., & Martinez-Trujillo, J. C. (2017). Neuronal population coding of perceived and memorized visual features in the lateral prefrontal cortex. Nature Communications, 8 (1), 1–13. doi:10.1038/ncomms15471

Miller, E. K., Erickson, C. A., & Desimone, R. (1996). Neural mechanisms of visual working memory in prefrontal cortex of the macaque. Journal of Neuroscience, 16, 5154–67. doi:10.1523/JNEUROSCI.16-16-05154.1996

Orbach J., & Fischer G. J. (1959). Bilateral resections of frontal granular cortex: Factors influencing delayed response and discrimination performance in monkeys. JAMA Neurology, 1(1), 78–86. doi:10.1001/archneur.1959.03840010080010

Petrides M. (2005). Lateral prefrontal cortex: Architectonic and functional organization. Philosophical transactions of the Royal Society of London. Series B, Biological sciences, 360 (1456), 781–795. doi:10.1098/rstb.2005.1631

Postle, B. R., Idzikowski, C., Sala, S. Della, Logie, R. H., & Baddeley, A. D. (1999). Quarterly Journal of Experimental Psychology, 59 (1), 100–120. doi:10.1080/17470210500151410

Roussy, M., Mendoza-Halliday, D., & Martinez-Trujillo, J. C. (2021a). Neural substrates of visual perception and working memory: Two sides of the same coin or two different coins? Frontiers in Neural Circuits, 15, 131. doi:10.3389/FNCIR.2021.764177/BIBTEX

Roussy, M., Luna, R., Duong, L., Corrigan, B., Gulli, R. A., Nogueira, R., … Martinez-Trujillo, J. C. (2021b). Ketamine disrupts naturalistic coding of working memory in primate lateral prefrontal cortex networks. Molecular Psychiatry, 1–16. doi:10.1038/s41380-021-01082-5

Sawaguchi, T., & Iba, M. (2001). Prefrontal cortical representation of visuospatial working memory in monkeys examined by local inactivation with muscimol. Journal of Neurophysiology, 86 (4), 2041–2053. doi:10.1152/jn.2001.86.4.2041

Sakagami, M., & Niki, H. (1994). Encoding of behavioral significance of visual stimuli by primate prefrontal neurons: Relation to relevant task conditions. Experimental Brain Research, 97 (3), 423–436. doi:10.1007/BF00241536

Suzuki, M., & Gottlieb, J. (2013). Distinct neural mechanisms of distractor suppression in the frontal and parietal lobe. Nature Neuroscience, 16 (1), 98–104. doi:10.1038/nn.3282

Takikawa, Y., Kawagoe, R., Itoh, H., Nakahara, H., & Hikosaka, O. (2002). Modulation of saccadic eye movements by predicted reward outcome. Experimental Brain Research, 142 (2), 284–291. doi:10.1007/s00221-001-0928-1

Quintana, J., Yajeya, J., Fuster, J. M. (1988). Prefrontal representation of stimulus attributes during delay tasks. I. Unit activity in cross-temporal integration of sensory and sensory-motor information. Brain Research, 474 (2), 211–221. doi:10.1016/0006-8993(88)90436-2

Yajeya, J., Quintana, J., & Fuster, J. M. (1988). Prefrontal representation of stimulus attributes during delay tasks. II. The role of behavioral significance. Brain Research, 474 (2), 222–230. doi:10.1016/0006-8993(88)90437-4

